# Multiplexed measurement of variant abundance and activity reveals VKOR topology, active site and human variant impact

**DOI:** 10.1101/2020.05.10.087312

**Authors:** Melissa A. Chiasson, Nathan J. Rollins, Jason J. Stephany, Katherine A. Sitko, Kenneth A. Matreyek, Marta Verby, Song Sun, Frederick P. Roth, Daniel DeSloover, Debora Marks, Allan E. Rettie, Douglas M. Fowler

## Abstract

Vitamin K epoxide reductase (VKOR) drives the vitamin K cycle, activating vitamin K-dependent blood clotting factors. VKOR is also the target of the widely used anticoagulant drug, warfarin Despite VKOR’s pivotal role in coagulation, its structure and active site remain poorly understood. In addition, VKOR variants can cause vitamin K-dependent clotting factor deficiency 2 or alter warfarin response. Here, we used multiplexed, sequencing-based assays to measure the effects of 2,695 VKOR missense variants on abundance and 697 variants on activity in cultured human cells. The large-scale functional data, along with an evolutionary coupling analysis, supports a four transmembrane domain topology, with variants in transmembrane domains exhibiting strongly deleterious effects on abundance and activity. Functionally constrained regions of the protein define the active site, and we find that, of four conserved cysteines putatively critical for function, only three are absolutely required. Finally, 25% of human VKOR missense variants show reduced abundance or activity, possibly conferring warfarin sensitivity or causing disease.

## INTRODUCTION

The enzyme vitamin K epoxide reductase (VKOR) drives the vitamin K cycle, which activates blood coagulation factors. VKOR, an endoplasmic reticulum (ER) localized transmembrane protein encoded by the gene *VKORC1*, reduces vitamin K quinone and vitamin K epoxide to vitamin K hydroquinone (Li et al., 2004; Rost et al., 2004). Vitamin K hydroquinone is required to enable gamma-glutamyl carboxylase (GGCX) to carboxylate Gla domains on vitamin K-dependent blood clotting factors. VKOR is inhibited by the anticoagulant drug warfarin (Czogalla et al., 2016; Zimmermann and Matschiner, 1974), and *VKORC1* polymorphisms contribute to an estimated ∼25% of warfarin dosing variability (Owen et al., 2010). For example, variation in *VKORC1* noncoding and coding sequence can cause warfarin resistance (weekly warfarin dose > 105 mg) or warfarin sensitivity (weekly warfarin dose < ∼10 mg) (Osinbowale et al., 2009; Yuan et al., 2005).

Though 15 million prescriptions are written for warfarin each year (https://www.clincalc.com), fundamental questions remain regarding its target, VKOR. For example, the structure of human VKOR is unsolved, though a bacterial homolog has been crystallized (Li et al., 2010). A homology model based on bacterial VKOR has four transmembrane domains, but the quality of the homology model is unclear, as human VKOR has only 12% sequence identity to bacterial VKOR. Moreover, experimental validation of VKOR topology yielded mixed results: similar biochemical assays suggested either three- or four-transmembrane-domain topologies (Schulman et al., 2010; Tie et al., 2012; Wu et al., 2018).

Topology informs basic aspects of VKOR function including where vitamin K and warfarin bind, so determining the correct topology and validating the homology model is critical. In particular, VKOR has four functionally important, absolutely conserved cysteines at positions 43, 51, 132, and 135, the orientation of which differs between the two proposed topologies. In the four transmembrane domain topology, all four cysteines are located on the ER lumenal side of the enzyme. In this topology, cysteines 43 and 51 are hypothesized to be “loop cysteines” that pass electrons from an ER-anchored reductase, possibly TMX (Schulman et al., 2010), to the active site (Rishavy et al., 2011). However, in the three transmembrane domain topology, these cysteines are located in the cytoplasm and other pathways would be required to convey electrons to the redox center. Even for non-catalytic residues, topology plays an important role. For example, vitamin K presumably binds near the redox center, and topology dictates which residues make up the substrate binding site.

To understand the effect of human variants and to define the vitamin K and warfarin binding sites, VKOR variant activity has been extensively studied in cell-based assays (Czogalla et al., 2016; Shen et al., 2017; Tie et al., 2013). In addition to activity, VKOR protein abundance has also been studied because abundance is an important driver of disease and warfarin response. For example, VKOR R98W is a decreased-abundance variant that, in homozygous carriers, causes vitamin K-dependent clotting factor deficiency 2 (Rost et al., 2004). A 5’ UTR polymorphism reduces VKOR abundance and can be used to predict warfarin sensitivity (Gong et al., 2011). However, so far, the activity and abundance of only a handful of VKOR variants has been tested.

Here, we used multiplexed, sequencing-based assays (Gasperini et al., 2016) to measure the effects of 2,695 VKOR missense variants on abundance and 697 variants on activity. Our analysis of the large-scale functional data supports a four transmembrane domain topology, which an orthogonal evolutionary coupling analysis confirmed. Next, we identified distinct mutational tolerance groups, which are concordant with a four transmembrane homology model. Combining this homology model with variant abundance and activity effects, we identified an active site that contains the catalytic residues C132 and C135 and shares six positions with a previously proposed vitamin K binding site (Czogalla et al., 2016). We found that of four conserved cysteines putatively critical for function, only three are absolutely required, and analyzed the mutational signatures of two putative ER retention motifs. Human *VKORC1* variants present in genetic databases and contributed by a commercial genetic testing laboratory were each classified based on abundance and activity. While most variants show wild type-like activity, 25% show low abundance or activity, which could confer warfarin sensitivity or cause disease in a homozygous context. Finally, we analyzed warfarin resistance variants and found that they span a range of abundances, indicating that increased abundance is an uncommon mechanism of warfarin resistance.

## RESULTS

### Multiplexed measurement of VKOR variant abundance using VAMP-seq

To measure the abundance of VKOR variants, we applied Variant Abundance by Massively Parallel sequencing (VAMP-seq), an assay we recently developed (Matreyek et al., 2018). In VAMP-seq, a protein variant is fused to eGFP with a short amino acid linker. If the variant is stable and properly folded, then the eGFP fusion will not be degraded, and cells will have high eGFP fluorescence. In contrast, if the variant causes the protein to misfold, protein quality control machinery will detect and degrade the eGFP fusion, leading to a decrease in eGFP signal (Fig. 1a). mCherry is also expressed from an internal ribosomal entry site (IRES) to control for expression. Differences in abundance are measured on a flow cytometer using the ratio of eGFP to mCherry signal. To determine whether VAMP-seq could be applied to VKOR, we fused eGFP to VKOR N- or C-terminally and found that both orientations had high eGFP signal (Figure 1—figure supplement 1). We compared N-terminally tagged wild type (WT) VKOR to R98W, a variant that ablates a putative ER retention motif and reduces abundance (Czogalla et al., 2014), and to TMD1Δ, a deletion of residues 10-30 which comprise the putative first transmembrane domain (TMD1; Fig. 1b). Both reduced-abundance variants exhibited much lower eGFP:mCherry ratios than WT, demonstrating that VAMP-seq could be applied to VKOR.

**Figure 1.**
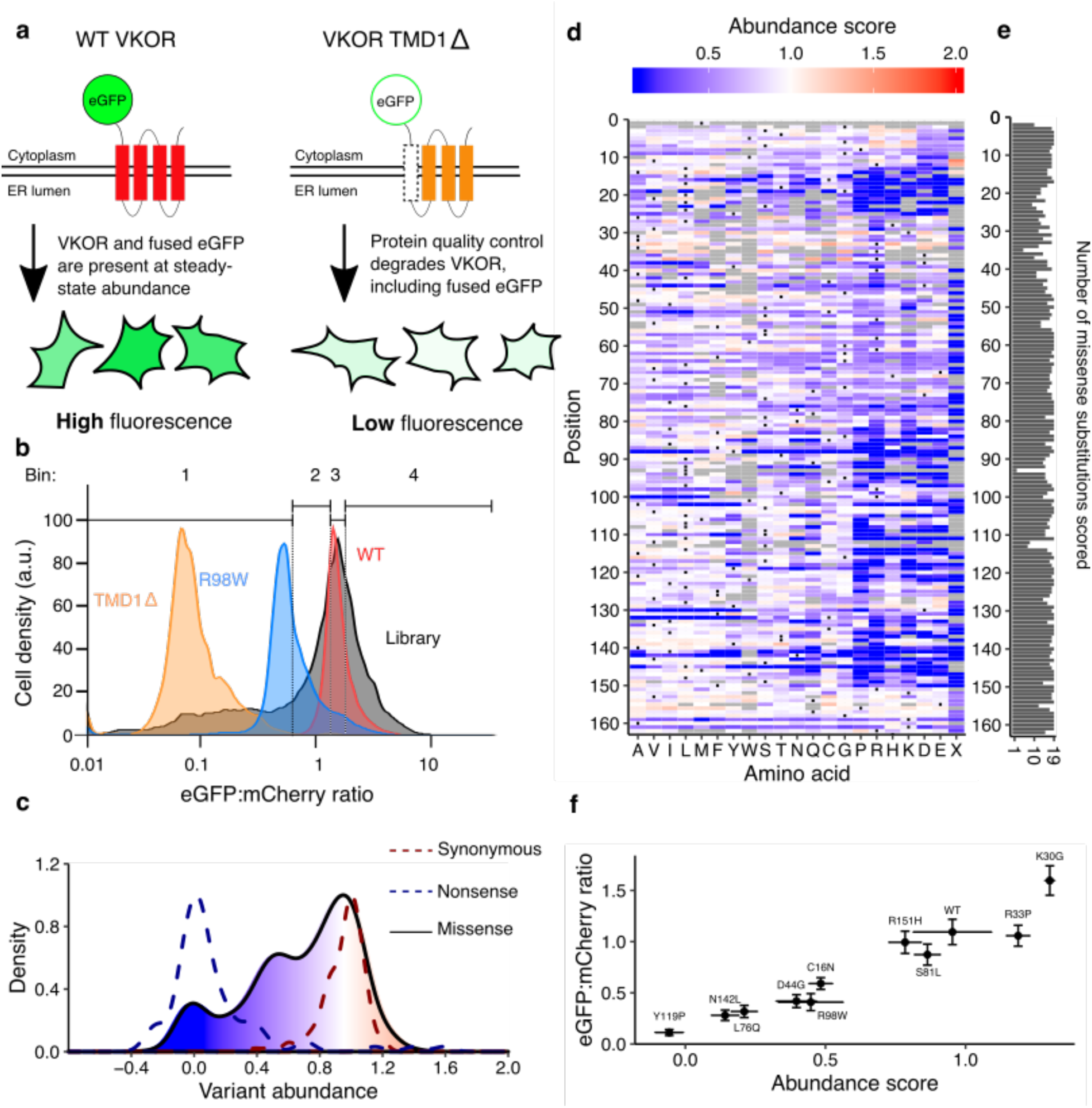
Multiplexed measurement of VKOR variant abundance using VAMP-seq. **a**, To measure abundance, an eGFP reporter is fused to VKOR. eGFP-tagged WT VKOR is folded correctly, leading to high eGFP fluorescence. However, a destabilized variant is degraded by protein quality control machinery, leading to low eGFP fluorescence. **b**, Flow cytometry is used to bin cells based on their eGFP:mCherry fluorescence intensity. Density plots of VKOR library expressing cells (grey, n = 12,109) relative to three controls: WT VKOR (red, n = 4,756), VKOR 98W (blue, n = 2,453), and VKOR TMD1Δ (orange, n = 2,204) are shown. Quartile bins for FACS of the library are marked. **c**, Abundance score density plots of nonsense variants (dashed blue line, n = 88), synonymous variants (dashed red line, n = 127), and missense variants (filled, solid line, n = 2,695). The missense variant density is colored as a gradient between the lowest 10% of abundance scores (blue), the WT abundance score (white) and abundance scores above WT (red). **d**, Heatmap showing abundance scores for each substitution at every position within VKOR. Heatmap color indicates abundance scores scaled as a gradient between the lowest 10% of abundance scores (blue), the WT abundance score (white), and abundance scores above WT (red). Grey bars indicate missing variants. Black dots indicate WT amino acids. **e**, Number of substitutions scored at each position for abundance. **f**, Scatterplot comparing VAMP-seq derived abundance scores to mean eGFP:mCherry (n = 1 replicate) ratios measured individually by flow cytometry. Variants were selected at random to span the abundance score range. **Figure 1-source data 1.** VKOR variant abundance and activity scores. **Figure 1-source data 2.** Flow cytometry for monoclonal validation of variants. 11 variants were run individually, values show mean and error for VAMP-seq score and eGFP:mcherry intensity.

We constructed a barcoded site-saturation mutagenesis VKOR library that covered 92.5% of all 3,240 possible missense variants. To express this library in HEK293T cells we used a Bxb1 recombinase landing pad system we previously developed (Matreyek et al., 2017). In this system, each cell expresses a single VKOR variant. Recombined, VKOR variant-expressing cells were then sorted into quartile bins based on their eGFP:mCherry ratios. Each bin was deeply sequenced, and abundance scores were calculated based on each variant’s distribution across bins. Raw abundance scores were normalized such that WT-like variants had a score of one and total loss of abundance variants had a score of zero (Fig. 1c). We performed seven replicates, which were well correlated (Figure 1—figure supplement 2, mean Pearson’s r = 0.73; mean Spearman’s ρ = 0.7, Supplementary Table 1). Abundance score means and confidence intervals for each variant were calculated from the replicates.

The final dataset describes the effect of 2,695 of the 3,240 possible missense VKOR variants on abundance (Fig. 1d and 1e). Validation of 10 randomly selected variants spanning the abundance score range showed high concordance between individual eGFP:mCherry ratios assessed by flow cytometry and VAMP-seq derived abundance scores (Fig. 1f, Pearson’s r = 0.96, Spearman’s ρ = 0.97).

### Multiplexed measurement of VKOR variant activity using a gamma-glutamyl carboxylation reporter

We also measured VKOR variant activity, adapting a HEK293 cell assay based on vitamin K-dependent gamma-glutamyl carboxylation of a cell-surface reporter protein (Haque et al., 2014). In this assay, if VKOR is active, a Factor IX domain reporter is carboxylated, secreted and retained on the cell surface where it is detected with a carboxylation-specific, fluorophore-labeled antibody. However, if VKOR is inactive, the reporter is not carboxylated and the antibody cannot bind (Fig. 2a). We modified the HEK293 activity reporter cell line to eliminate endogenous VKOR activity by knocking out both *VKORC1* and its paralog, VKORC1-like 1 (*VKORC1L1*) (Tie et al., 2013) (Figure 2—figure supplement 1). We also installed a Bxb1 landing pad to facilitate expression of individual VKOR variants or libraries (Figure 2—figure supplement 1). Recombination of WT *VKORC1* into the landing pad of the HEK293 VKOR activity reporter cell line yielded robust reporter activation, demonstrating that the reporter line could be used to assess the activity of a library of VKOR variants (Fig. 2b).

**Figure 2.**
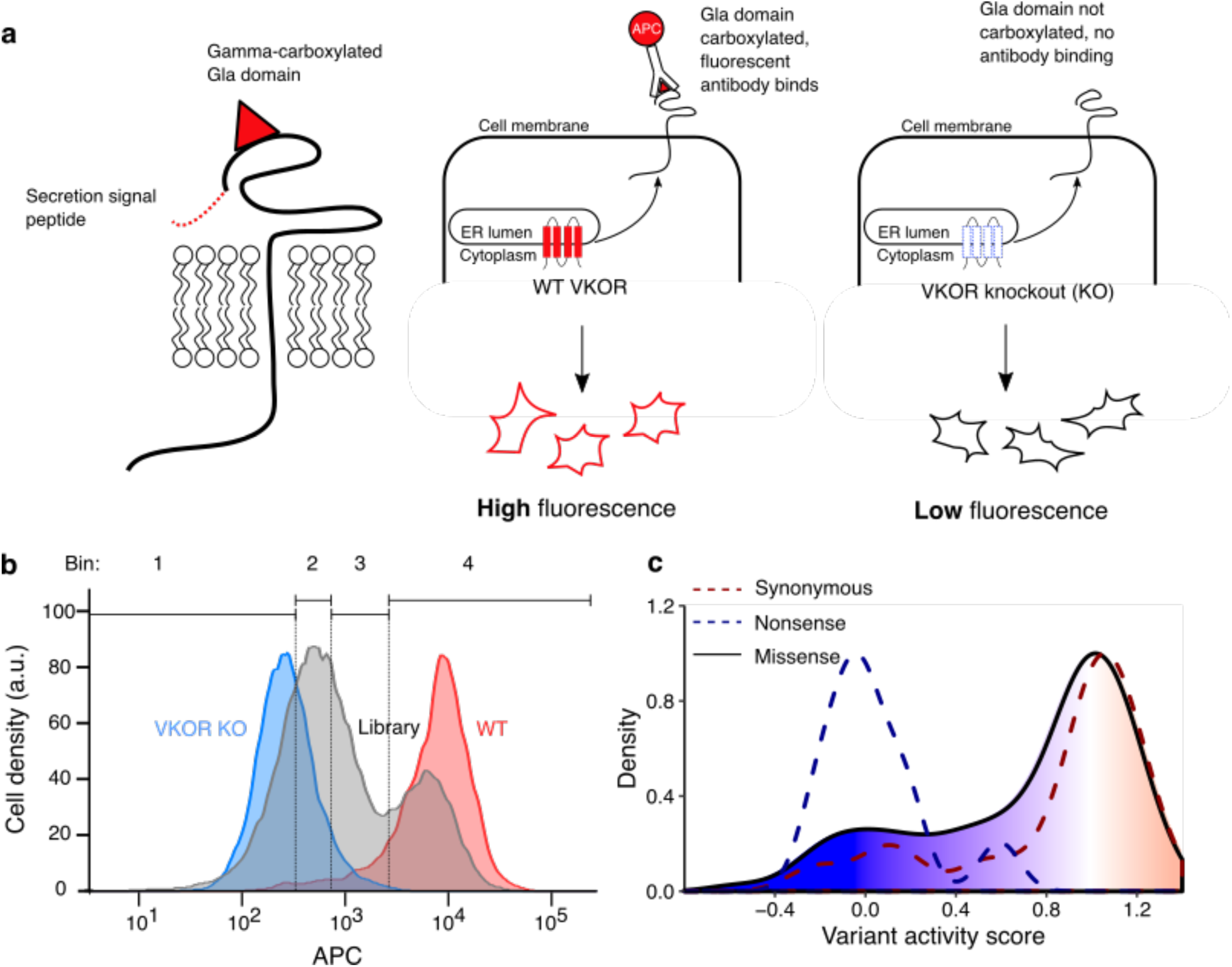
Multiplexed measurement of VKOR variant activity using a gamma-glutamyl carboxylation reporter. **a, left panel**, a Factor IX Gla domain reporter is expressed inHEK293 cells and consists of a prothrombin pre-pro-peptide which allows for processing and secretion, a Factor IX Gla domain, and Proline rich Gla protein 2 (PRGP2) transmembrane and cytoplasmic domains. **middle panel,** Cells expressing WT VKOR carboxylate the reporter Gla domain, which, upon trafficking to the cell surface, can be stained using a carboxylation-specific antibody conjugated to the fluorophore APC. VKOR knockout cells do not carboxylate the reporter, so the fluorescent antibody does not bind. **b**, Density plots of HEK293 activity reporter cells stained with APC-labeled carboxylation-specific antibody expressing no VKOR (blue, n = 7,188), WT VKOR (red, n = 4,107), or the VKOR variant library (grey, n = 41,418). Quartile bins for FACS of the library are marked. **c**, Activity score density plots of nonsense variants (dashed blue line, n = 14), synonymous variants (dashed red line, n = 35), and missense variants (filled, solid line, n = 697). The missense variant density is colored as a gradient between the lowest 10% of activity scores (blue), the WT activity score (white) and activity scores above WT (red).

We recombined a library of *VKORC1* variants into the HEK293 activity reporter cell line, and sorted recombinant cells into quartile bins based on carboxylation-specific antibody binding. Each bin was deeply sequenced and, as for VAMP-seq, an activity score was computed for each variant. Final activity scores and confidence intervals were computed from six replicates for a total of 697 missense variants, 21.5% of those possible (Figure 2—figure supplement 2, mean Pearson’s r = 0.62 and mean Spearman’s ρ = 0.56, Supplementary Table 2). Our activity score density plot showed that most variants had WT-like activity scores (Fig. 2c).

### Human VKOR has four transmembrane domains

Two different domain models, one with three transmembrane domains and another with four, have been proposed for human VKOR (Li et al., 2010; Tie et al., 2012) (Fig. 3a). Because charged amino acids occur infrequently in transmembrane domains and should be less tolerated, we reasoned we could discriminate between these two models using a sliding window average of the effect of charged substitutions on VKOR abundance (A. Elazar et al., 2016; Sharpe et al., 2010). We found four clearly demarcated regions where charged substitutions profoundly reduced VKOR abundance, relative to aliphatic substitutions (Fig. 3b). To exclude the possibility that the eGFP tag used in our VAMP-seq assay somehow affected topology, we also analyzed the activity score data. The activity data, derived using native, untagged VKOR, revealed the same four minima as the abundance data (Fig. 3c). In addition to these four minima, we also observed an activity score minimum at position 57, corresponding to a conserved serine at this position. This serine occurs at the end of the lumenal half-helix hypothesized to shield the active site from non-specific oxidation, so it is likely this signal is the result of disruption of that half helix. Together, these results strongly support the hypothesis that, like its distant bacterial homolog, human VKOR has four transmembrane domains.

**Figure 3.**
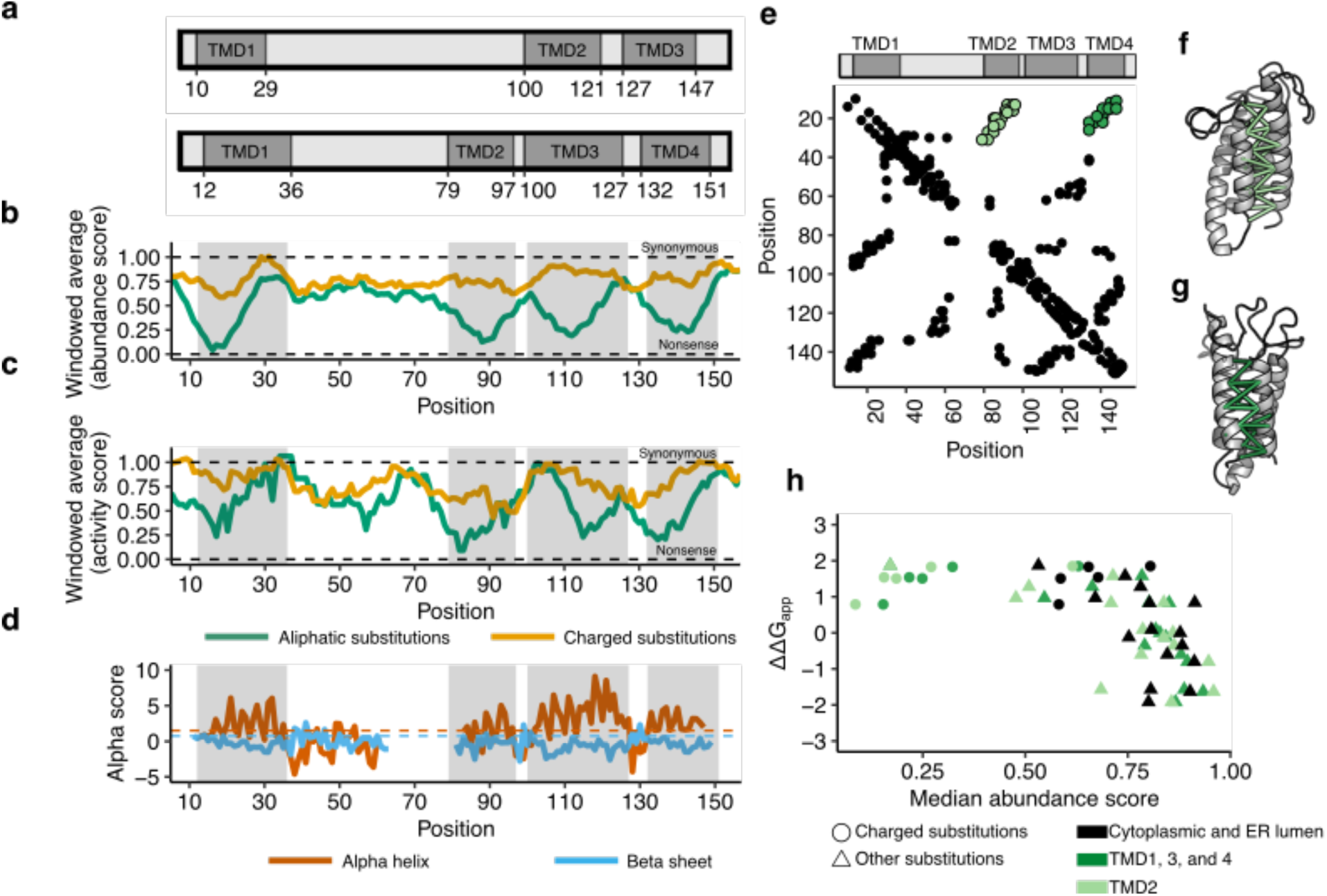
Abundance, activity, and evolutionary data support four transmembrane domains. **a**, Three and four transmembrane domain (TMD) models of VKOR, with TMDs in dark grey (Li et al., 2010; Tie et al., 2012). **b**, Windowed abundance score means (width = 10 positions) for charged substitutions (green) and aliphatic substitutions (gold). Dark grey boxes correspond to TMDs proposed in the four domain model. Dashed lines show median synonymous and the nonsense abundance scores. **c**, Windowed activity score means (width = 10 positions) for charged substitutions (green) and aliphatic substitutions (gold). Boxes and dashed lines as described in **b**. **d**, Secondary structure classification from local evolutionary couplings shown as alpha scores calculated for alpha helices (red) and beta sheets (blue). Dashed lines show significance cut-offs for alpha helices (1.5, red) and beta sheets (0.75, blue) (Toth-Petroczy et al., 2016). **e**, A contact map derived from evolutionary couplings. Black points show pairs of positions with significant coupling. Light green points show predicted contacts between TMD1 and TMD2. Dark green points show predicted contacts between TMD1 and TMD4. **f**, Predicted tertiary contacts between TMD1-TMD2 (shown in light green in **e**) and **g**, TMD1-TMD4 (shown in dark green in **e**) shown on the evolutionary couplings-derived hVKOR structural model. **h**, Scatterplot comparing change in free energy for membrane insertion (A. A. Elazar et al., 2016) (ΔΔG_app_) to median abundance score for each amino acid substitution. Cytoplasmic and lumenal positions shown in black, TMD2 in light green, and TMDs 1, 3, and 4 in dark green. Charged substitutions shown as circles, all other substitutions as triangles. **Figure 3-source data 1.** Evolutionary couplings secondary structure predictions. Rows show position, with columns showing alpha helix or beta sheet values and predictions. **Figure 3-source data 2.** Evolutionary couplings 3D contact predictions. Rows show pairs of residues with contact probabilities. **Figure 3-source data 3.** Insertion energies from Elazar et al., 2016. Amino acids with calculated insertion energy.

To validate these findings, we performed evolutionary coupling analysis to infer the three-dimensional structure suggested by co-evolution. We aligned 2,770 VKOR sequences from both eukaryotes and prokaryotes and identified coupled residues using the EVcouplings software (Hopf et al., 2012; Marks et al., 2011). Local patterns of evolutionary couplings (i.e. between nearby positions, *i* to *i*+4) supported a four-helix topology. The helices predicted by these local evolutionary couplings overlapped 70 of the 82 residues in alpha-helices of the bacterial structure (PDB 4NV5) (Shen et al., 2017) and included in our alignment, non-gapped in >70% of aligned VKOR sequences (hyper-geometric test p-value = 3.26^-23^, Fig. 3d).

We identified non-local evolutionary coupling patterns characteristic of three-dimensional contacts, which also strongly supported the four transmembrane domain model. Using these contacts, we computationally folded human VKOR, yielding a modeled structure similar to the bacterial structure (RMSD = 2.58 ÅA over 97/143 C_alpha_, Figure 3—figure supplement 1). The predicted tertiary structure hasd a four-helix topology, with antiparallel contacts between transmembrane domains 1 and 2 (Fig. 3e, Fig. 3f) and between transmembrane domains 1 and 4 (Fig. 3e, Fig. 3g), which are only possible in a four-helix topology.

Comparison of our abundance data to the energy required to insert different amino acids into the membrane yields additional evidence for the four transmembrane domain model. The apparent change in free energy (ΔΔG_app_) of insertion relative to wild type for every amino acid has been determined experimentally using deep mutational scanning of bacterial membrane proteins (A. A. Elazar et al., 2016). Median abundance score and ΔΔG_app_ for each amino acid are correlated (Fig. 3h). In particular, the large energetic cost of insertion of transmembrane domains with charged amino acids is apparent, including within the second transmembrane domain TMD2. Beyond insertion energies of individual amino acids, the overall hydrophobicity of transmembrane helices contributes to membrane protein insertion (A. A. Elazar et al., 2016), as well as topology (A. Elazar et al., 2016) and degradation (Guerriero et al., 2017). To determine whether overall helix hydrophobicity was a large factor contributing to abundance scores, we calculated the free energy for insertion (ΔG_helix_) of each helix in the four transmembrane domain model using the ΔG prediction server v1.0 (Hessa et al., 2007). The four helices of VKOR have different ΔG_helix_, with only transmembrane domain 3 having favorable ΔG_helix_ for insertion (TMD1: 0.435, TMD2: 1.551, TMD3: −1.749, and TMD4: 1.734). Interestingly, we observed that TMD3 has a high density of substitutions with WT-like scores (Figure 3—figure supplement 2), suggesting that TMD3’s favorable insertion energy might explain its mutational tolerance.

### Detailed structural context of VKOR variant abundance effects

Having confirmed that human VKOR has four transmembrane domains, we next explored the detailed pattern of mutational effects we observed in the context of a four transmembrane domain homology model. We generated a homology model of human VKOR with I-TASSER using the bacterial VKOR structure (Shen et al., 2017; Yang et al., 2015). We performed hierarchical clustering of positions based on abundance scores, which yielded four groups of positions with characteristic mutational patterns (Fig. 4a). In Group 1, most substitutions were neutral or increased abundance; in Group 2, charged amino acid and proline substitutions decreased abundance; in Group 3, all substitutions decreased abundance; and in Group 4, all substitutions decreased abundance profoundly. Each group corresponded to a spatially distinct region of the homology model structure (Fig. 4b).

**Figure 4.**
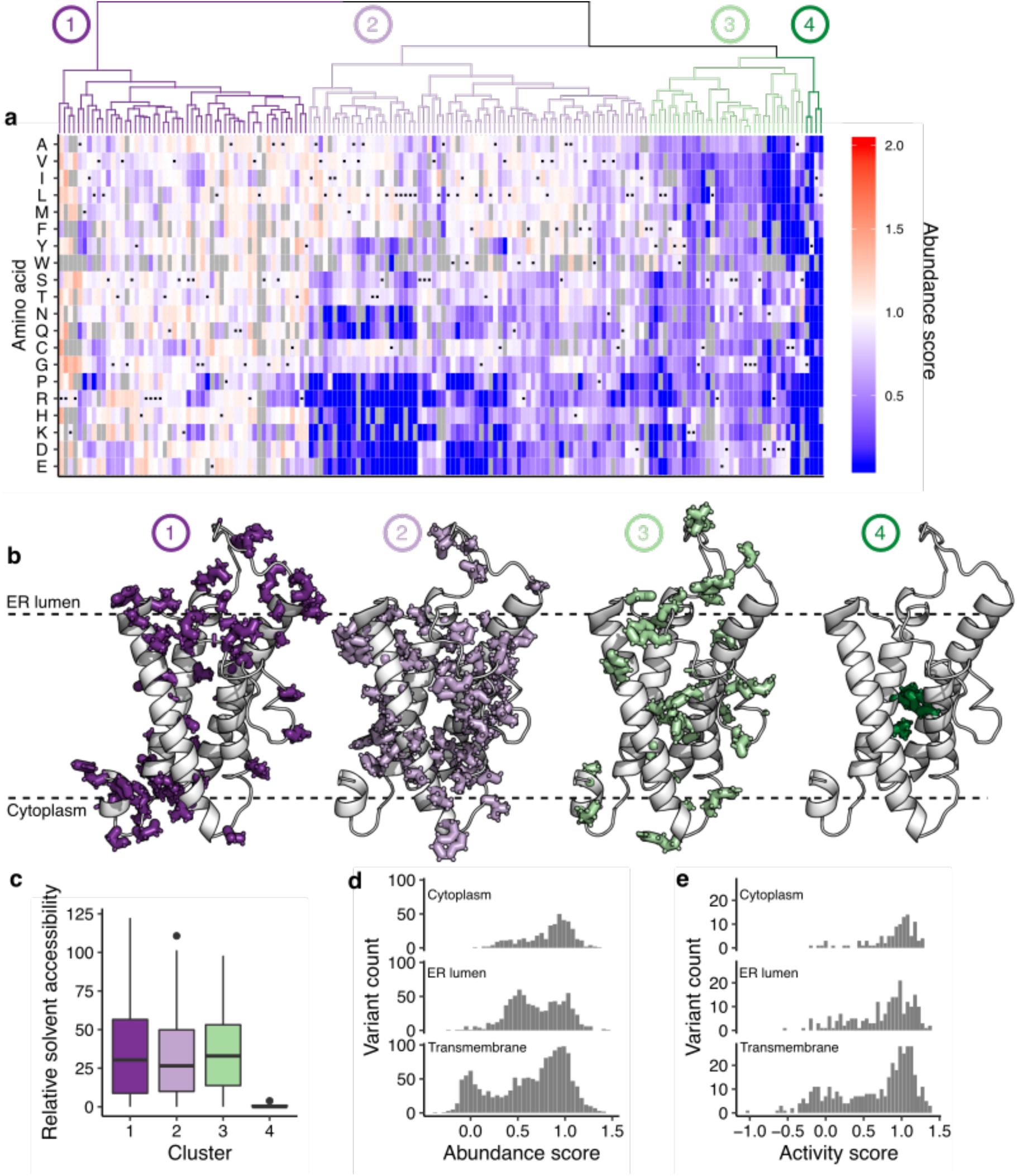
Hierarchical clustering of abundance scores and distributions of abundance and activity scores by domain. **a**, A heatmap showing hierarchical clustering of positions based on abundance score vectors, with the dendrogram above. Groups of positions, chosen based on the dendrogram, are numbered and colored. Heatmap color indicates abundance scores scaled as a gradient between the lowest 10% of abundance scores (blue), the WT abundance score (white) and abundance scores above WT (red). Grey bars indicate missing variants. Black dots indicate WT amino acids. **b**, Positions in groups 1-4 shown on the VKOR homology model, with numbers and colors corresponding to panel **a**. **c**, Boxplot showing relative solvent accessibility of positions in each cluster determined using DSSP (Kabsch and Sander, 1983; Touw et al., 2015) and colored as in **b**. Bold black line shows median, box shows 25th and 75th percentile. Line shows 1.5 interquartile range above and below percentiles, and outliers are shown as black points. **d**, Histograms of abundance scores for missense variants in the cytoplasmic, ER lumenal, or transmembrane domains. **e**, Histograms of activity scores for missense variants in the cytoplasmic, ER lumenal, or transmembrane domains.

Group 1 positions were located in or adjacent to cytoplasmic and ER lumenal loops, which were more tolerant of substitutions than the transmembrane domains. At four Group 1 positions, K30, R33, R35, and R37, almost every substitution increased abundance. These positively charged positions are positioned either at the edge of TMD1 (K30) or in the ER lumen directly abutting the top of TMD1 (R33, R35, and R37). The “positive inside rule” (von Heijne, 1989), suggests that positive charges in membrane proteins generally reside in the cytoplasm, and this phenomenon is important for driving topology and membrane insertion (A. Elazar et al., 2016; Nilsson and von Heijne, 1990; von Heijne, 1989). K30, R33, R35, and R37 violate the positive inside rule, and substitutions at these positions may increase abundance by reducing charge inside the ER, reducing topological frustration or increasing membrane insertion efficiency. Compared to the other 12 arginine and lysine positions in WT VKOR, K30, R33, R35, and R37 are the only ones where substitutions generally increased abundance (Figure 4—figure supplement 1). Our observations are consistent with a screen of rat VKOR variants intended to improve protein expression in *E. coli* where deletion of positions 31 to 33 increased protein levels (Hatahet et al., 2015).

In Group 2, charged amino acids or proline substitutions generally decreased abundance. Group 2 consisted mostly of transmembrane positions that had side chains projecting into the lipid bilayer. Such transmembrane positions usually have hydrophobic, nonpolar side chains (Ulmschneider and Sansom, 2001). Proline has poor helix forming propensity, explaining why proline substitutions decreased abundance at these positions. Group 3 consisted of a mixture of cytoplasmic, ER lumenal and transmembrane positions where most substitutions decreased abundance. The cytoplasmic positions in this group included the putative dilysine ER localization motif at positions 159 and 161. Also in this group were R98, part of another putative ER retention motif at positions 98 and 100, and a glycine adjacent to TMD1 at position nine. The transmembrane positions had side chains projecting towards neighboring transmembrane helices, suggesting that, as for other membrane proteins (Fleming and Engelman, 2001; Mravic et al., 2019), intramolecular sidechain packing is important for abundance.

Finally, substitutions in Group 4, consisting of positions G19, Y88, I141, and L145, resulted in catastrophic loss of abundance. These positions are all in transmembrane domains with side chains projecting into the interior of the protein. On the basis of strict mutational intolerance of these positions, we hypothesized that their coordinated side chain packing comprises the core of the VKOR four helix bundle. Indeed, Group 4 residues had dramatically lower relative solvent accessibility than Groups 1-3 (Fig. 4c).

The four transmembrane domain homology models also allowed us to explain VKOR’s unusual trimodal distribution of variant abundance scores. Previous VAMP-seq derived abundance score distributions for the cytosolic proteins TPMT and PTEN were bimodal (Figure 4—figure supplement 2) (Matreyek et al., 2018), and 15 of 16 deep mutational scans of other soluble proteins using a variety of other assays also exhibited bimodal functional score distributions (Gray et al., 2017). Because VKOR is an ER resident, transmembrane protein, we hypothesized that its unusual trimodal abundance score distribution resulted from transmembrane domain substitutions. Indeed, the lowest mode of the distribution was composed almost exclusively of deleterious transmembrane domain substitutions (Fig. 4d). In contrast, the intermediate mode consisted of substitutions in the ER lumen, cytoplasm, and transmembrane domains. Similarly, substitutions that profoundly decreased activity occurred in transmembrane domains (Fig. 4e).

### Variant activity and abundance identify functionally constrained regions of VKOR

We reasoned that our activity and abundance data could reveal the location of functionally important positions in VKOR, including the active site, since functionally important positions should have many loss-of-activity but few loss-of-abundance variants. Thus, we calculated the specific activity for each variant by taking the ratio of its rescaled activity score and abundance scores (see Methods) ratio. We computed the median specific activity for each position; substitutions at positions with low median specific activity generally have low activity relative to their abundance. We set a specific activity threshold based on two absolutely conserved cysteines that form VKOR’s redox center, C132 and C135. Using this threshold, positions with the lowest 12.5% of specific activity scores and with at least four variants scored for activity were deemed functionally constrained and mapped on the homology model of VKOR (Fig. 5a, Figure 5—figure supplement 1). These 11 functionally constrained positions are organized around C132 and C135 and define, at least in part, the VKOR active site (Fig. 5b,c, Figure 5—figure supplements 1). Among the functionally constrained positions are six positions previously identified in vitamin K docking simulations (Czogalla et al., 2016) (Figure 5—figure supplement 1), including F55, which is hypothesized to bind vitamin K. Three functionally constrained positions, G60, R61, and A121, did not match any position in the active site predicted by docking, but were immediate neighbors of W59 and L120, positions that were.

**Figure 5.**
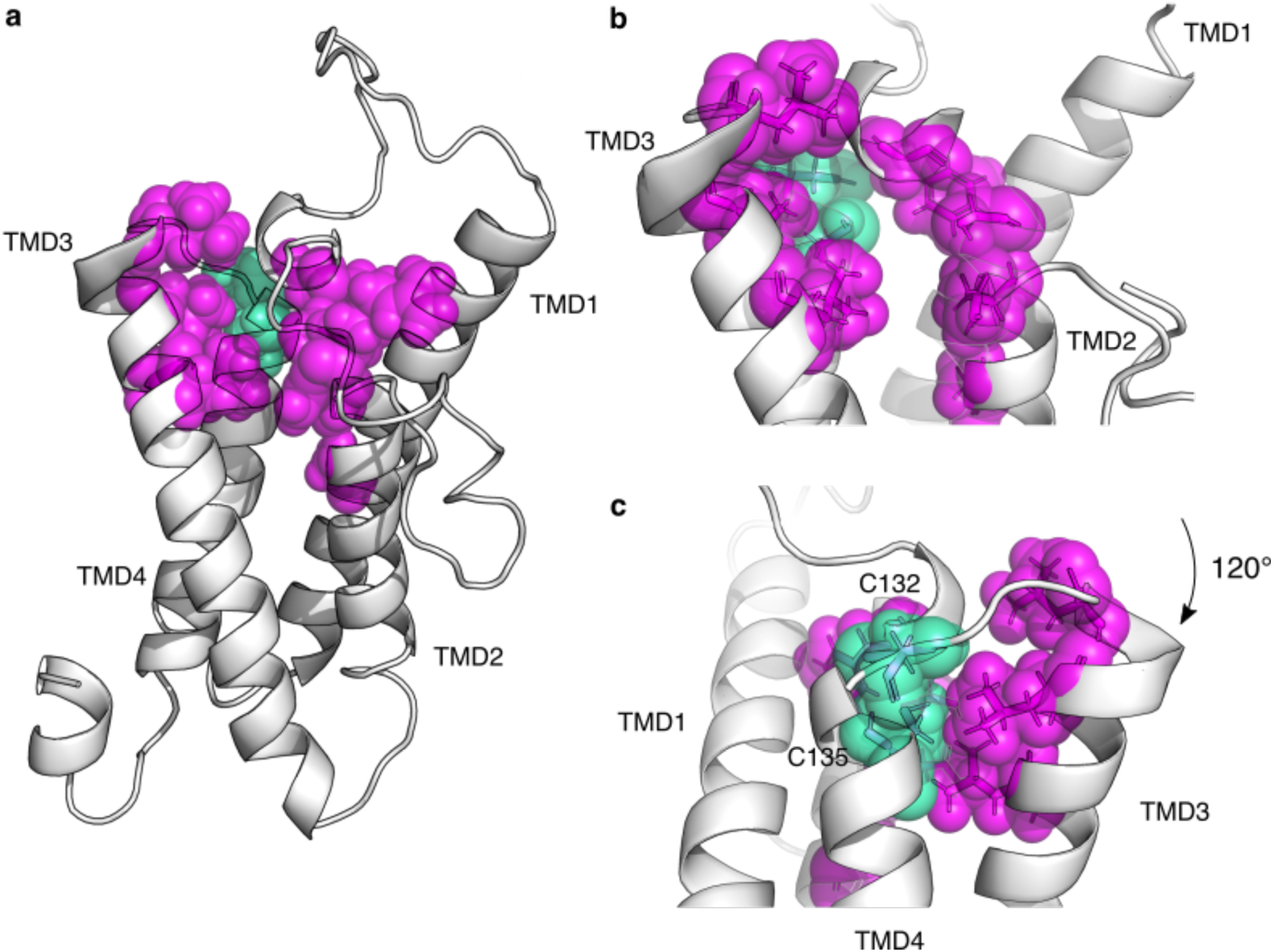
Functionally constrained positions reveal VKOR active site and critical cysteines. **a**, Positions with the lowest 12.5% of median specific activity scores and at least four variants scored for activity are shown as magenta spheres on the VKOR homology model. Cysteines C132, and C135, also in the bottom 12.5% of median specific activity scores, are shown in green spheres. **b**, Magnified view of the redox center cysteines (positions 132, and 135, green spheres) and surrounding residues that define the active site (magenta spheres). Residues shown in transparent spheres, with side chains also shown in sticks. **c**, panel **b** rotated 120°C. **Figure 5-source data 1.** VKOR positional abundance and activity scores. Rows show positions, with columns showing median abundance score, median activity score, rescaled scores, and specific activity score.

Besides C132 and C135, VKOR has two additional absolutely conserved cysteines, C43 and C51. In the four transmembrane domain model, C43 and C51 are postulated to be loop cysteines that relay electrons to the C132/C135 redox center (Liu et al., 2014). We classified C43 as having low specific activity, but we only observed one variant at this position, so it was not included in our set of functionally constrained positions (Figure 5—figure supplement 2). In contrast, substitutions at C51 resulted in only modest activity loss, a phenomenon that has been observed previously (Shen et al., 2017). Interestingly, every substitution at C51 and 15 of 19 at C132 decreased VKOR abundance (Figure 5—figure supplement 2). Inside cells, the majority of VKOR molecules have a C51-C132 disulfide bond, and warfarin binds to this redox state of VKOR (Shen et al., 2017). Since disruption of this disulfide bond apparently impacts abundance as well as activity, this bond may be important for VKOR folding and stability.

VKOR is thought to contain two sequences important for ER localization. The first is a diarginine motif (RxR) at positions 98-100, and the second is a dilysine motif (KXKXX) at positions 159-163. While we did not directly measure localization, we found that only six of 19 R98 variants and seven of 14 R100 variants resulted in low abundance (Figure 5—figure supplement 3). In contrast, nearly all variants at K159 (14 of 18) and K161 (17 of 19) resulted in low abundance (Figure 5—figure supplement 3). A histidine substitution was tolerated at position 161, which mimics the KXHXX motif commonly found in coronaviruses and a small number of human proteins (Ma and Goldberg, 2013). Because protein localization and degradation are coupled (Hessa et al., 2011), we suggest that the reductions in abundance we observe are the result of degradation caused by mislocalization, and that the dilysine motif at positions 159-163 is essential for VKOR ER localization. Overall, comparison of VKOR variant activity and abundance revealed functionally important regions, refining our understanding of the active site, redox-active cysteines, and ER retention motifs.

### Functional consequences of VKOR variants observed in humans

Variation in VKOR is linked to both disease and warfarin response, but the overwhelming majority of VKOR variants found in humans so far have unknown effects. Thus, we curated a total of 215 variants that had either been previously reported in the literature as affecting warfarin response (Supplementary Table 3), were in ClinVar (Landrum et al., 2014), were in gnomAD v2 or v3 (Karczewski et al., 2019), or were present in individuals whose healthcare provider had ordered a multi-gene panel test from a commercial testing laboratory (Color Genomics) (Supplementary Table 4). Of eight variants present in ClinVar, we included only one (D36Y) in our analysis as it was the only variant reviewed by an expert panel (Kurnik et al., 2012). 159 variants were present in gnomAD, and all but one missense variant (D36Y) had population frequencies less than 0.2%. 28 were literature-curated warfarin response variants, only 12 variants of which were in one of the databases surveyed. D36Y was the only warfarin response variant present in all databases, ClinVar, gnomAD, and Color (Figure 6—figure supplement 1).

We classified 193 of the 215 variants we curated according to their abundance (Supplementary Table 4). All synonymous variants with the exception of two were WT-like or possibly WT-like, while the three nonsense variants scored had as having low abundance (Fig. 6a). Missense variants spanned all abundance categories, with 129 (60%) having WT-like or possibly WT-like abundance. 30 missense variants were low abundance, and 12 were high abundance. The single known pathogenic variant R98W was low abundance (Fig. 6b). We also classified 54 variants according to their activity (Supplementary Table 4). Only one variant, A115V, exhibited low activity. It had WT-like abundance, indicating that the loss of activity is not due to loss of abundance.

**Figure 6.**
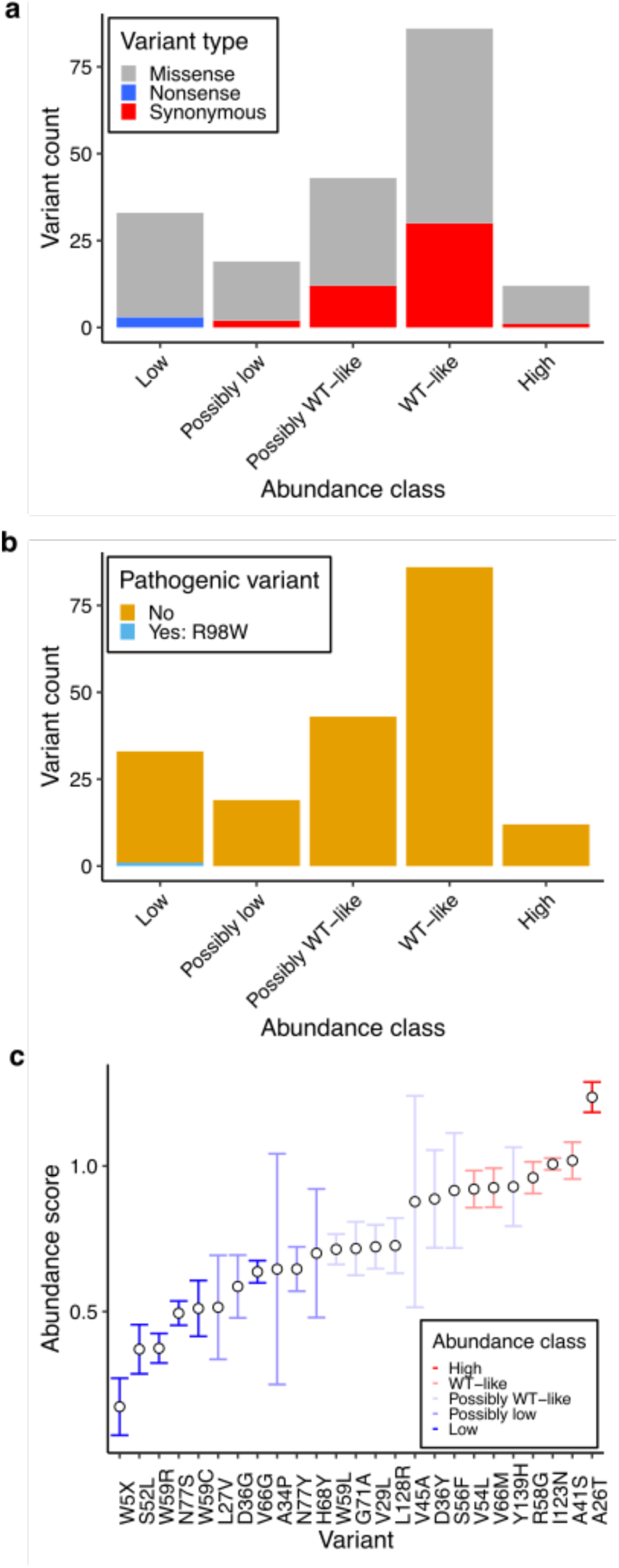
Characterization of human variants using abundance and activity data. **a**, Histogram of abundance classifications for variants from gnomAD, ClinVar, and Color Genomics. Nonsense variants colored in blue, synonymous in red, and missense in grey. **b**, Histogram of abundance classifications for same variants in **a**, colored by pathogenicity. The only variant known to cause disease, R98W, is colored in blue. All other variants shown in yellow. **c**, Scatterplot showing abundance scores for literature-curated warfarin resistance variants. Bars show standard error and are colored by abundance class. Variants are arranged in order of abundance score.

We examined warfarin response variants including W5X, the only variant observed so far linked to human warfarin sensitivity (Oldenburg et al., 2004). As expected, W5X was low abundance, reinforcing that heterozygous loss of VKOR is the cause of warfarin sensitivity in carriers of this variant. Warfarin resistance variants, on the other hand, are predicted to abrogate warfarin binding (Li et al., 2010), but it is unclear whether these variants have appreciable effects on abundance or activity. We found that warfarin resistance variants span a range of abundances and that the distribution of warfarin resistant variant abundance was not different from missense variants generally (Fig. 6c, two-sided Kolmogorov-Smirnov test p= 0.438). Five warfarin-resistance variants had low abundance, suggesting that these variants must block drug binding or increase activity to confer resistance. One variant, A26T, had high abundance, a possible mechanism of warfarin resistance. The five warfarin resistance variants, R58G, W59L, V66M, G71A, and N77S, whose activity we scored, were all WT-like. Thus, our abundance and activity data are consistent with warfarin resistance arising largely from variants that block warfarin binding.

## DISCUSSION

We conducted multiplexed assays to measure the effects of 2,695 VKOR variants on abundance and 697 variants on activity. Both abundance and activity data provided evidence for a four transmembrane topology, which was further supported by evolutionary couplings analysis. We evaluated a VKOR homology model in the context of the patterns of variant effects on abundance we measured, and found that the homology model could explain these patterns. Low specific activity residues mapped onto this homology model identify, at least in part, the active site, which largely overlaps with the results of a vitamin K docking simulation (Czogalla et al., 2016). Our active site is shallower than what the docking simulation predicts; this is the result of low abundance scores at some of the deeper, transmembrane positions predicted by docking to bind the isoprenoid chain of vitamin K (F87, Y88), and poor coverage of activity scores for other positions (V112, S113). In light of the fact that substitutions at F87 and Y88 resulted in low abundance, we note that the modeled vitamin K binding mode would disrupt packing of VKOR core residues and require repacking of helices to maintain protein stability (Merkle et al., 2018). In addition to the active site, substitutions at the dilysine and, to a lesser extent, the diarginine ER localization motifs caused abundance loss.

We also used our large-scale functional data to analyze 215 VKOR variants found in humans. 16% of these variants affect neither activity nor abundance; we identified 54 previously uncharacterized low abundance or low activity variants that could be pathogenic or alter warfarin response. We found that only one warfarin resistance variant had increased abundance, indicating that increased abundance is not a pervasive warfarin resistance mechanism. All five of the warfarin resistance variants whose activity we scored were WT-like. Taken together these data support the notion that warfarin resistance generally involves alterations to warfarin binding rather than abundance or activity. We analyzed one known warfarin sensitivity variant, W5X, and found that it is low abundance, suggesting that one should not exclude the possibility that any of the 52 other low abundance variants, if found in a person, also confer warfarin sensitivity.

While our VKOR variant abundance and activity data illuminates various aspects of VKOR’s structure and function, the data have limitations. For example, neither assay captures variant effects on mRNA splicing. Both assays have limited dynamic ranges, meaning that subtle effects on abundance or activity cannot be discerned. In addition, both assays have inherent noise, largely arising from the limited number of cells we can sample due to the bottleneck of cell sorting. We account for this noise by filtering each dataset based on variant frequency and presenting a confidence interval for each abundance and activity score.

In the future, we envision that the assays we used could be employed to better understand VKOR’s interaction with warfarin. Here, we could measure warfarin’s effect on both variant abundance and activity, mapping the warfarin binding site more finely. In addition, we could identify warfarin resistance mutations that have not yet been observed in the clinic and group variants by their putative resistance mechanism. Overall, our work highlights the value of multiplexed assays of variant effect for better understanding protein structure, function and human variant effects.

---

**Figure 1—figure supplement 1.**
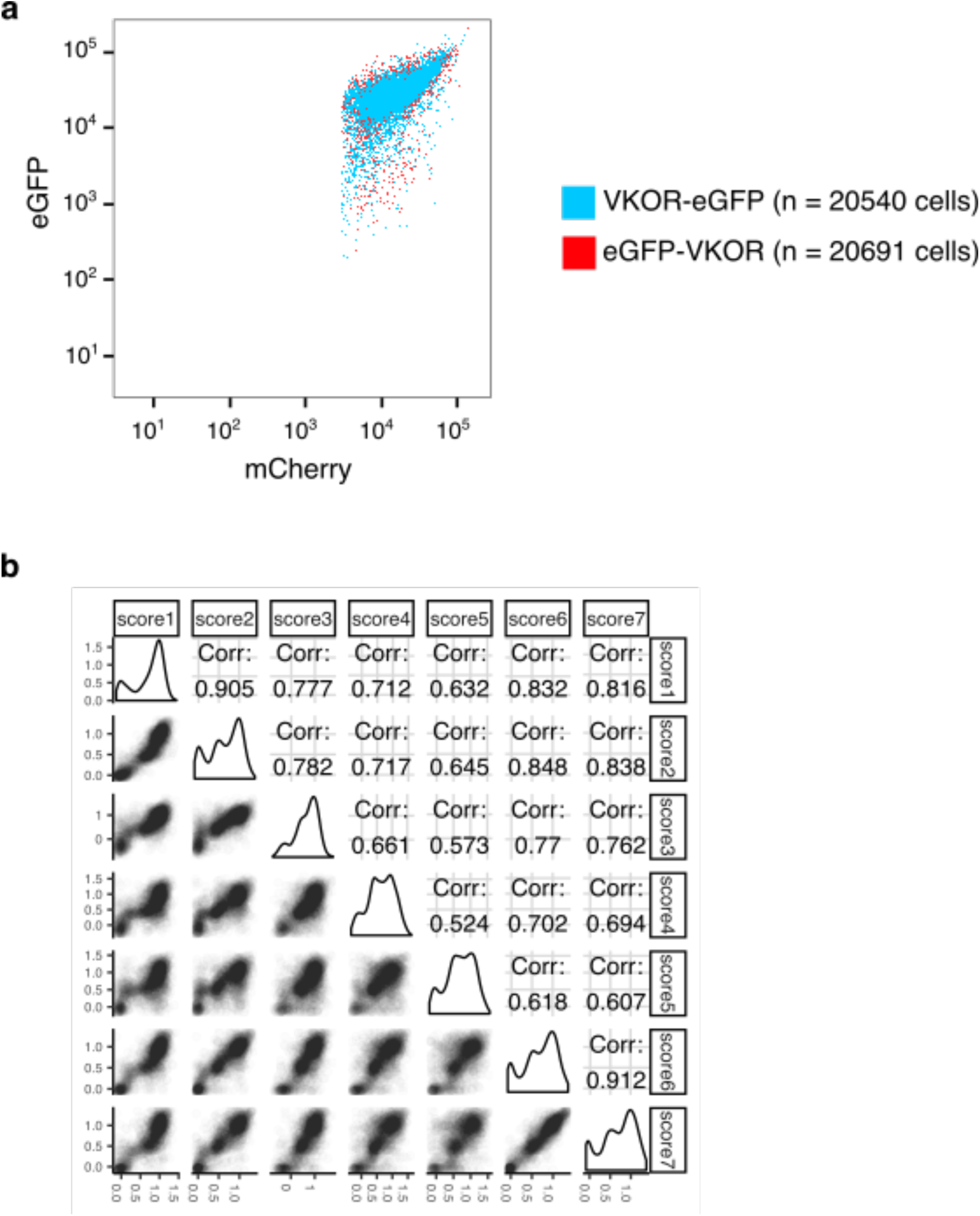
VKOR abundance assay pilot experiment and replicate correlations. **a**, Scatterplot of eGFP vs. mCherry fluorescence for cells expressing either C-terminally eGFP-tagged VKOR (VKOR-eGFP, blue) or N-terminally eGFP-tagged VKOR (eGFP-VKOR, red). **b**, Pairwise abundance score correlations between replicate sorting experiments. Seven VAMP-seq replicates were performed. Pearson’s correlation coefficients are shown. Score numbers in this figure correspond to replicate numbers shown in Supplementary Table 1.

**Figure 2—figure supplement 1.**
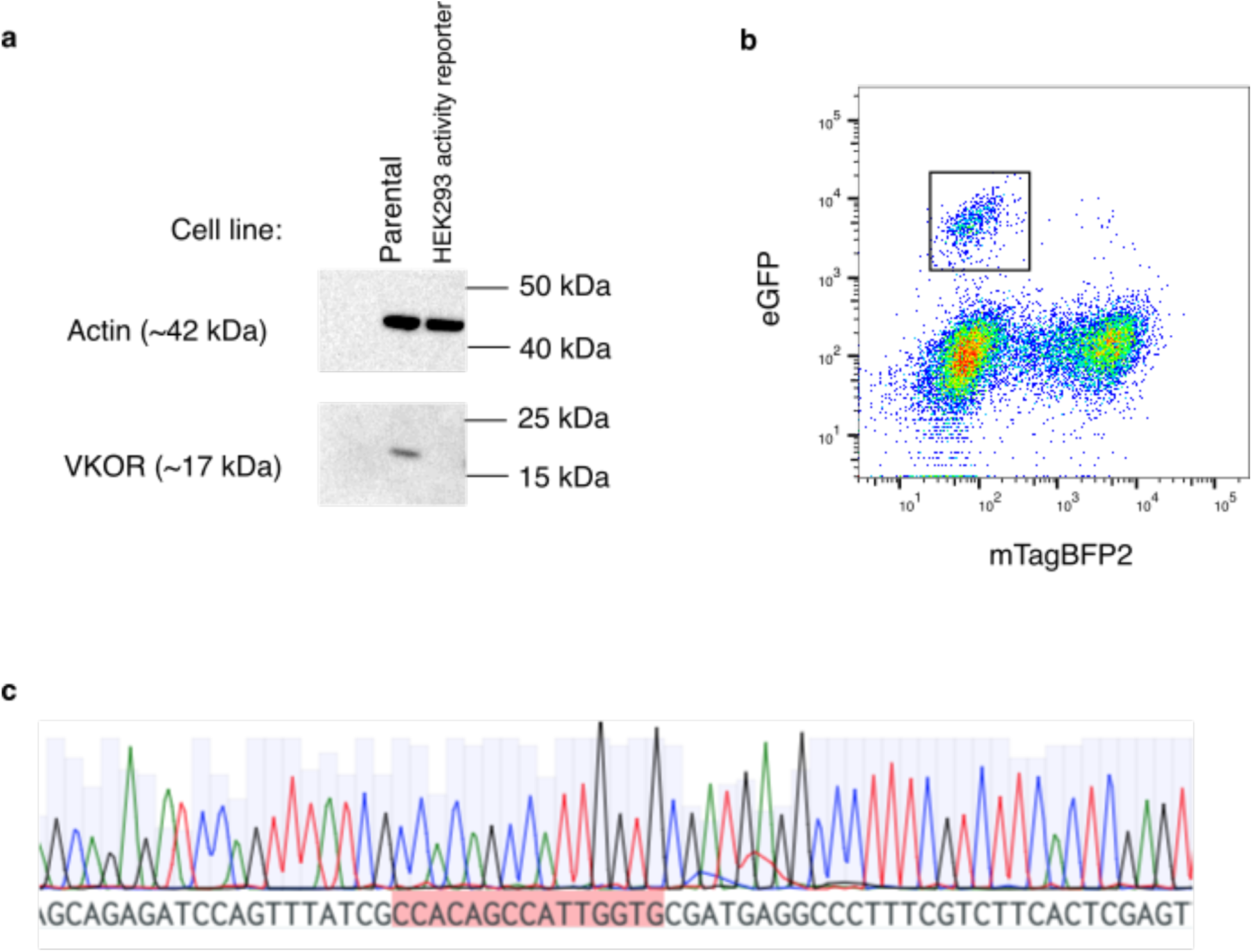
HEK293 VKOR activity reporter cell line characterization. **a**, Western blot of parental cell line vs. HEK293 activity reporter cell line. Loading control is actin (42 kDa). VKOR was probed using an antibody generated against a peptide from the C-terminal of VKOR (FRKVQEPQGKAKRH)(Hallgren et al., 2006). The band for VKOR at 17 kDA is visible in the parental cell line but is not present in the HEK293 activity reporter cell line. **b**, Scatterplot showing mTagBFP2 vs. eGFP mean fluorescence intensities for HEK293 activity reporter cells recombined with a construct encoding WT VKOR followed by internal ribosomal entry sequence and eGFP. The emergence of a distinct recombined population that is eGFP positive and mTagBFP2 negative (black outline, n = 768 cells) supports the presence of a single landing pad into the cell genome, and not multiple insertions. **c**, A chromatogram showing the barcode sequence of the landing pad inserted at the *AAVS1* locus in the HEK293 activity reporter cell line. The presence of a single barcode, highlighted in red, instead of mixed peaks, supports insertion of one landing pad rather than multiple landing pads.

**Figure 2—figure supplement 2.**
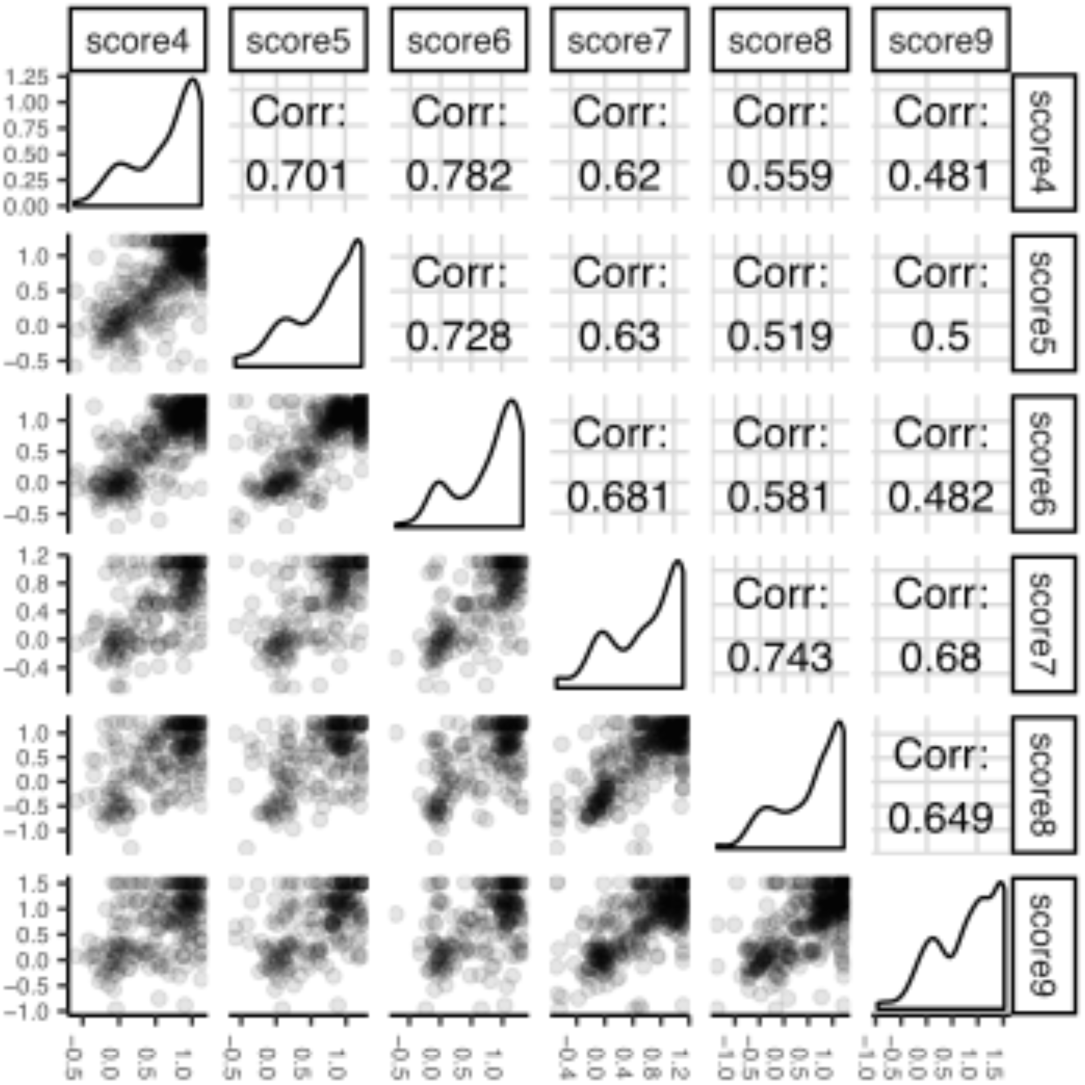
Correlations of activity assay replicates. Pairwise score correlations between replicate sorting experiments of VKOR activity. Six replicates of the activity assay were performed. Pearson’s correlation coefficients are shown. Score numbers in this panel correspond to replicate numbers shown in Supplementary Table 2.

**Figure 3—figure supplement 1.**
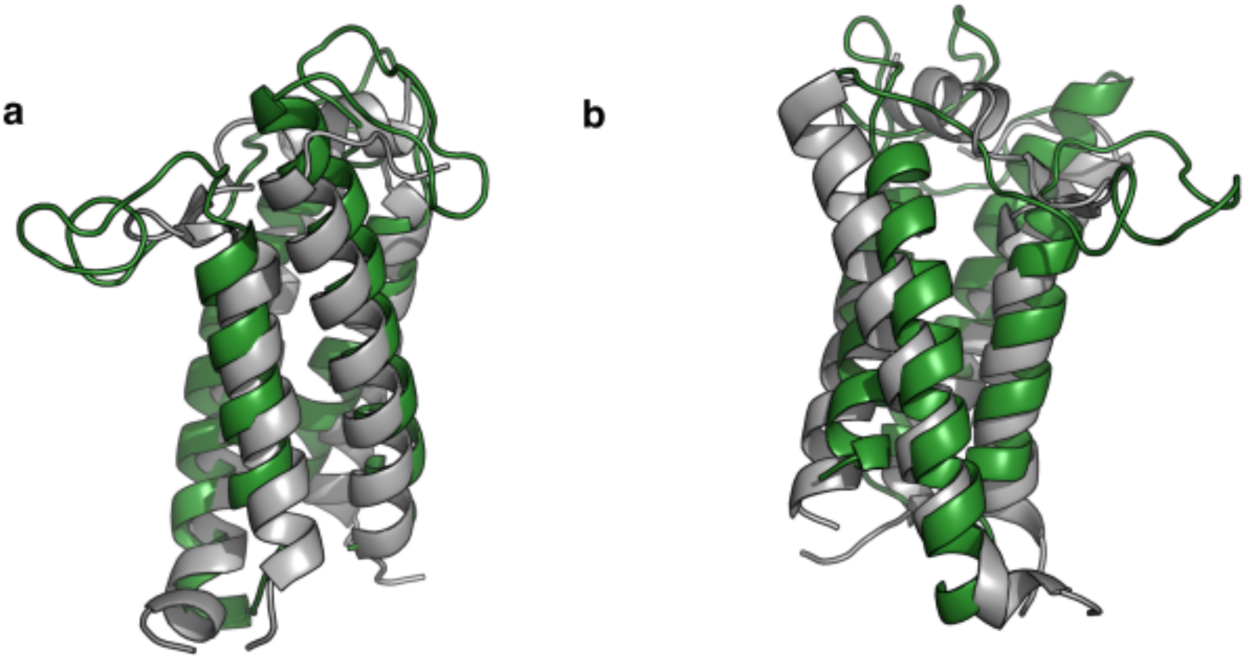
Bacterial VKOR structure and EV-couplings folded model are highly similar. **a**, Pymol graphic showing overlap between EVcouplings-folded model of VKOR (shown as a cartoon in green) compared to the bacterial structure (PDB: 4NV5, shown as a cartoon in grey). **b**, shows the same two structures, rotated 120°C.

**Figure 3—figure supplement 2.**
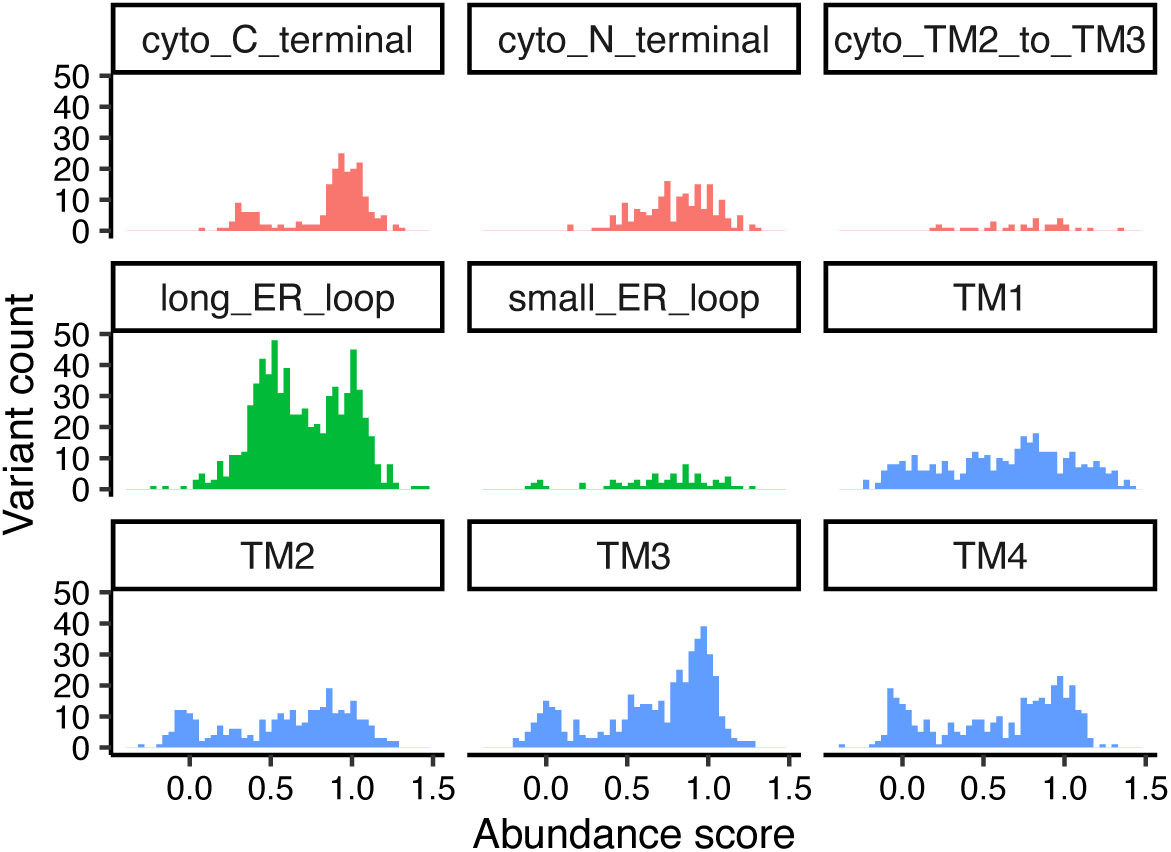
Specific domain abundance scores. Histograms of abundance scores for missense variants, grouped by domain and colored by cytoplasmic, ER lumenal, or transmembrane localization.

**Figure 4—figure supplement 1.**
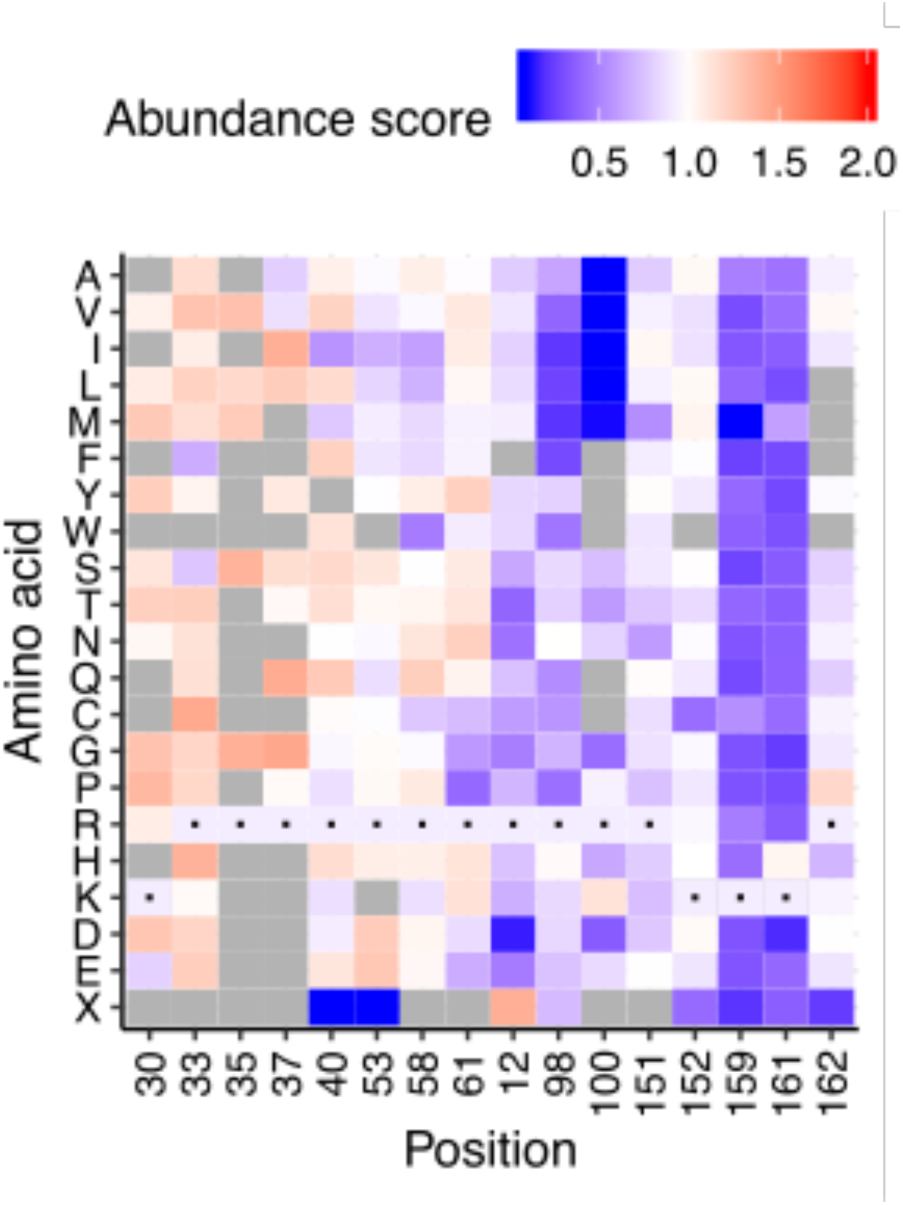
TMD1-adjacent positive residues show pattern of increased abundance. Heatmap of abundance scores for all arginines and lysines in VKOR. First four positions (K30, K33, K35, K37) are in or proximal to transmembrane domain 1. Heatmap color indicates abundance scores scaled as a gradient between the lowest 10% of abundance scores (blue), the WT abundance score (white) and abundance scores above WT (red). Grey bars indicate missing variants. Black dots indicate WT amino acids.

**Figure 4—figure supplement 2.**
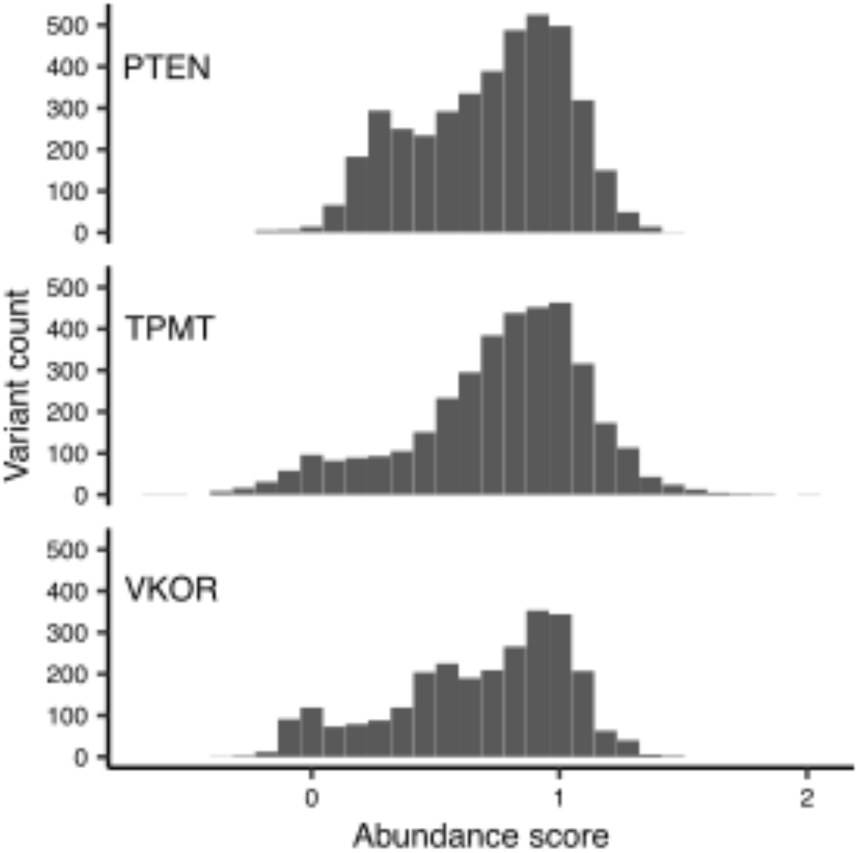
Trimodality of missense variant abundance scores is unique to VKOR. Histograms of abundance scores for missense variants for three proteins: PTEN, TPMT, and VKOR.

**Figure 5—figure supplement 1.**
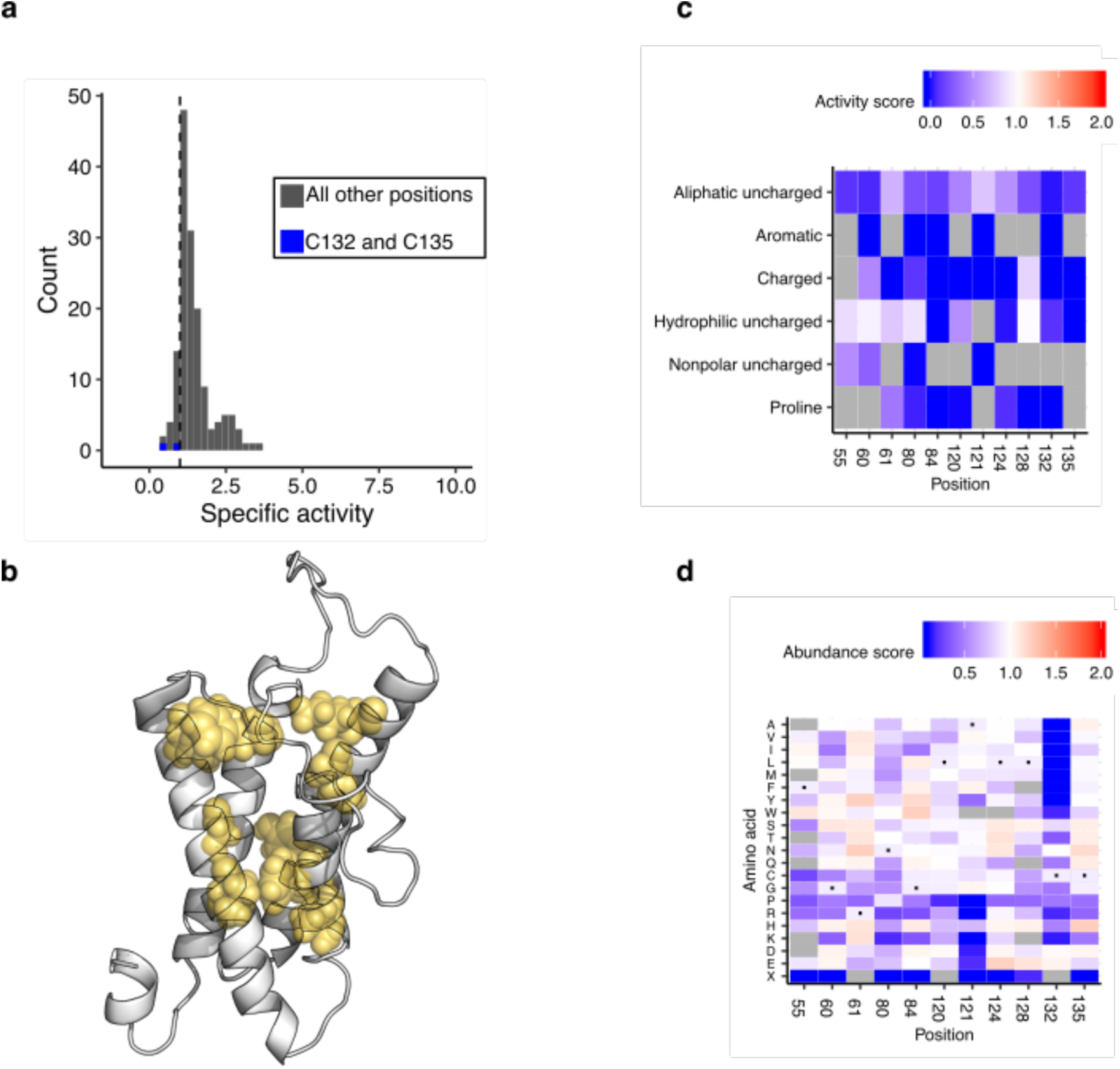
VKOR active site analysis. Histogram of specific activity, with catalytic cysteines C132 and C135 labeled in blue. Dashed line demarcates bottom 12.5%. **b**, Active site positions as defined by computational docking, shown on the homology model as yellow spheres(Czogalla et al., 2016). **c**, Heatmap of activity scores for residues with lowest 12.5% of specific activity scores, collapsed by amino acid class. Color indicates activity scores scaled as a gradient between the lowest 10% of activity scores (blue), the WT activity score (white) and activity scores above WT (red). Grey indicates missing data. **d**, Heatmap of abundance scores for residues with lowest 12.5% of specific activity scores. Color legend same as described in **c,** applied to abundance scores.

**Figure 5—figure supplement 2.**
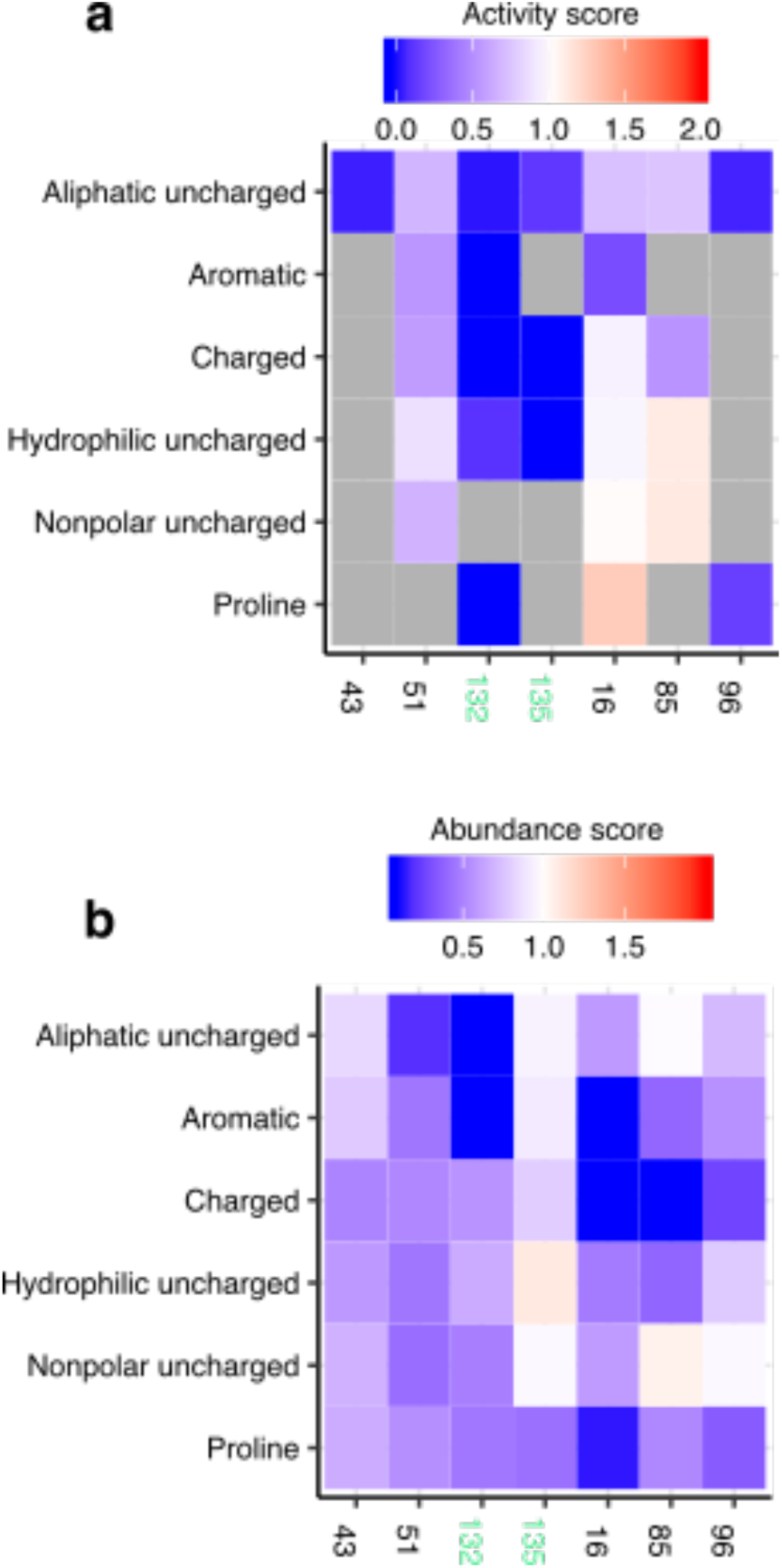
Conserved cysteine analysis. **a**, Heatmap of activity scores for cysteines. Catalytic cysteines C132 and C135 labeled in green. Color indicates activity scores scaled as a gradient between the lowest 10% of activity scores (blue), the WT activity score (white) and activity scores above WT (red). Grey indicates missing data. **b**, Heatmap of abundance scores for cysteines. Catalytic cysteines C132 and C135 labeled in green. Color legend same as described in **a,** applied to abundance scores.

**Figure 5—figure supplement 3.**
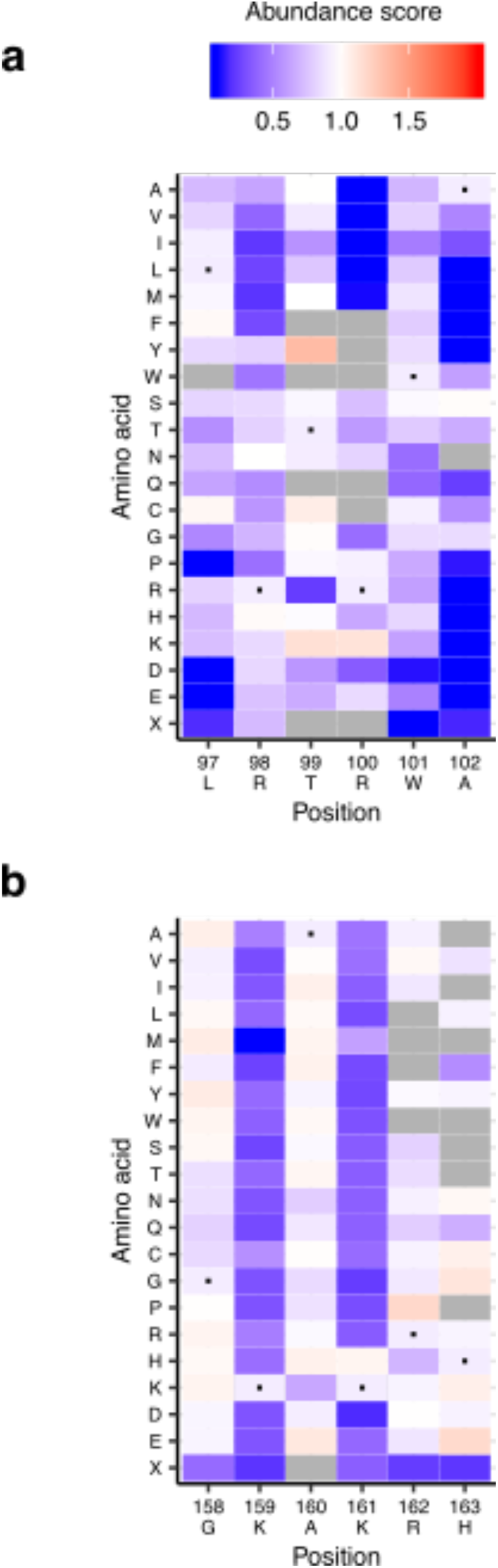
VKOR localization motif analysis. **a**, Heatmap of abundance scores for diarginine ER retention motif. X-axis shows residues and position. Color indicates abundance scores scaled as a gradient between the lowest 10% of abundance scores (blue), the WT abundance score (white) and abundance scores above WT (red). Grey indicates missing data. **b**, Heatmap of abundance score for dilysine ER retention motif. X-axis shows residues and position.

**Figure 6—figure supplement 1.**
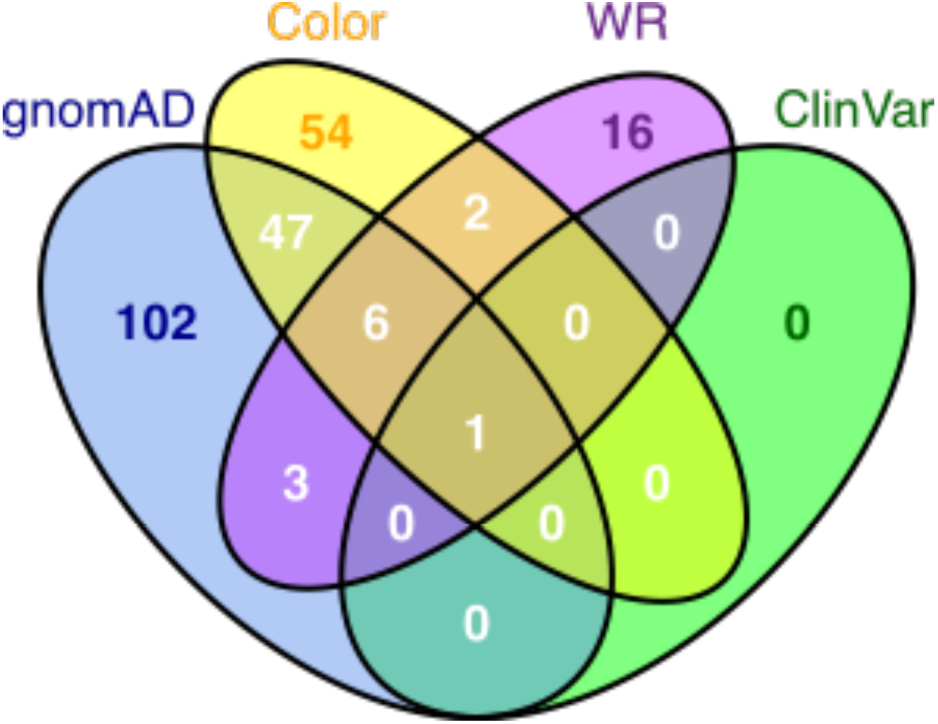
Human VKOR variant curation summary. Venn diagram of VKOR missense variants present in gnomAD v2 and v3, ClinVar, Color Genomics, a commercial genetic testing company, and literature-reported warfarin resistant variants.

**Supplementary Table 1.** The seven replicates of VAMP-seq performed with cells recombined and sorted for each.

| Replicate | Cells recombined | Cells sorted in four-way sort |
| --- | --- | --- |
| 1 | 256,324 | 200,000 |
| 2 | 256,324 | 200,000 |
| 3 | 111,570 | 183,000 |
| 4 | 155,169 | 100,000 |
| 5 | 105,000 | 103,000 |
| 6 | 54,045 | 120,000 |
| 7 | 54,045 | 125,000 |

**Supplementary Table 2.** The six replicates of the activity assay performed with cells recombined and sorted for each.

| Replicate | Cells recombined | Cells sorted in four-way sort |
| --- | --- | --- |
| 1 | 85,492 | 200,000 |
| 2 | 85,492 | 150,000 |
| 3 | 85,492 | 100,000 |
| 4 | 165,000 | 90,000 |
| 5 | 165,000 | 90,000 |
| 6 | 165,000 | 100,000 |

**Supplementary Table 3.**
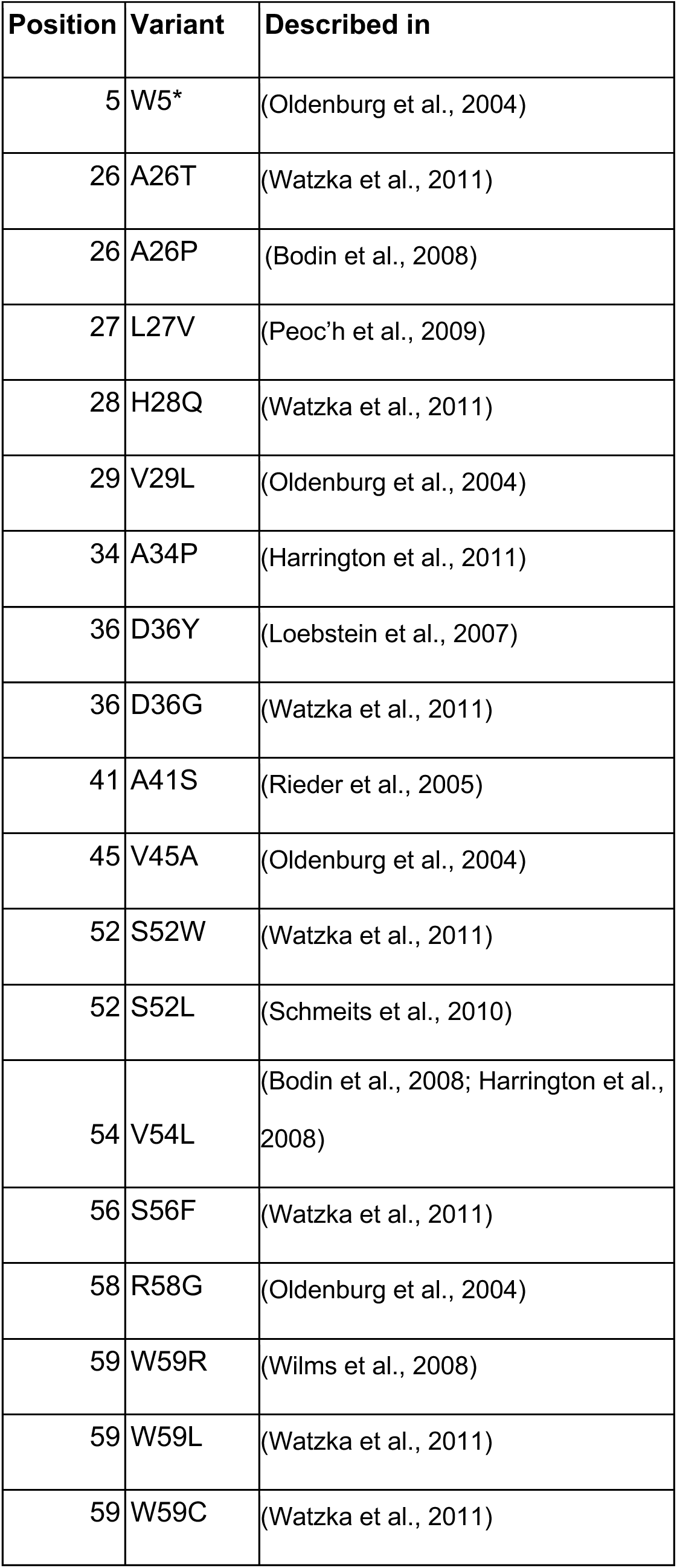
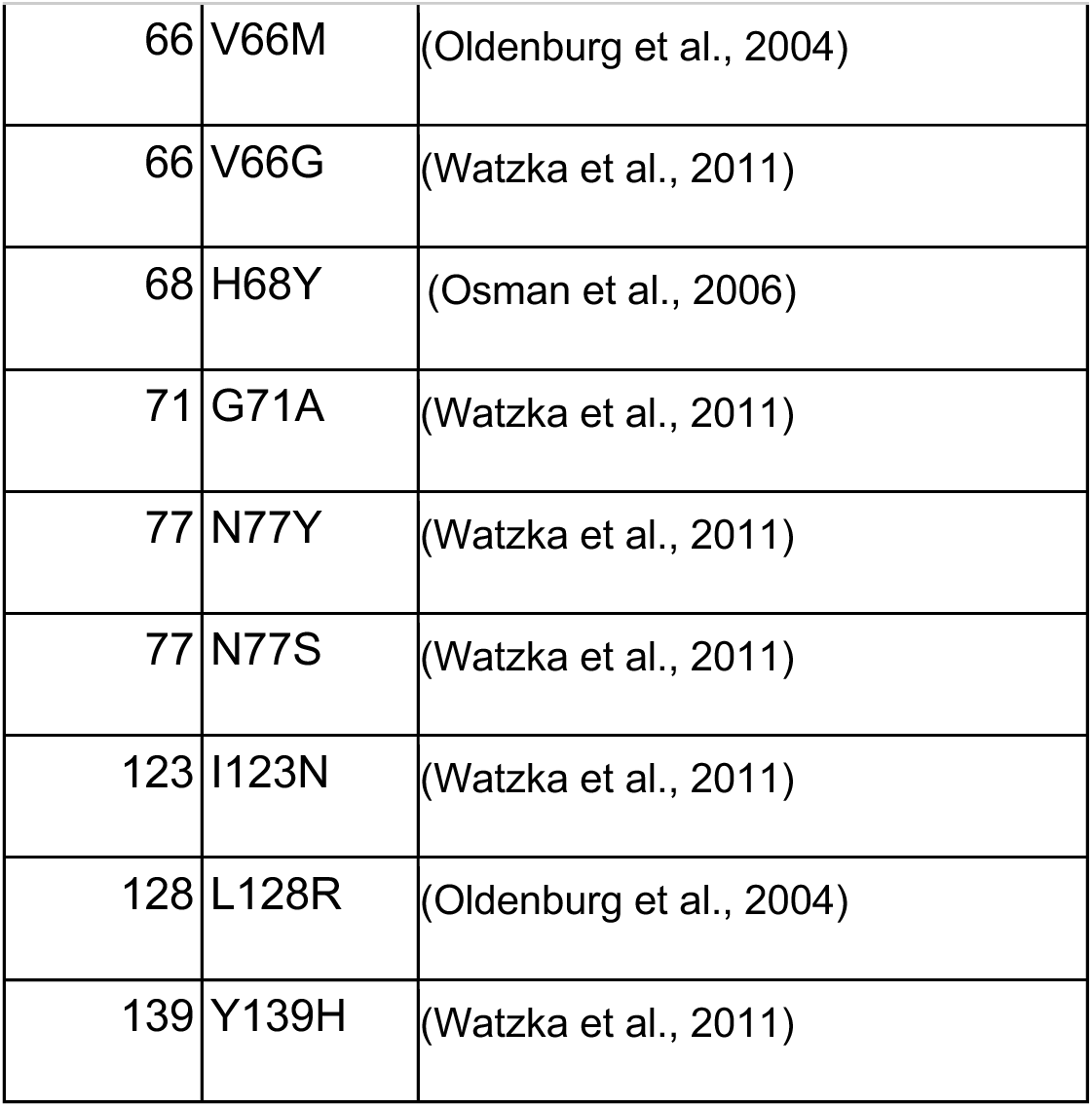
Variants found in humans that cause warfarin sensitivity or resistance, and references in which they were first reported.

| Position | Variant | Described in |
| --- | --- | --- |
| 5 | W5* | (Oldenburg et al., 2004) |
| 26 | A26T | (Watzka et al., 2011) |
| 26 | A26P | (Bodin et al., 2008) |
| 27 | L27V | (Peoc'h et al., 2009) |
| 28 | H28Q | (Watzka et al., 2011) |
| 29 | V29L | (Oldenburg et al., 2004) |
| 34 | A34P | (Harrington et al., 2011) |
| 36 | D36Y | (Loebstein et al., 2007) |
| 36 | D36G | (Watzka et al., 2011) |
| 41 | A41S | (Rieder et al., 2005) |
| 45 | V45A | (Oldenburg et al., 2004) |
| 52 | S52W | (Watzka et al., 2011) |
| 52 | S52L | (Schmeits et al., 2010) |
| 54 | V54L | (Bodin et al., 2008; Harrington et al., 2008) |
| 56 | S56F | (Watzka et al., 2011) |
| 58 | R58G | (Oldenburg et al., 2004) |
| 59 | W59R | (Wilms et al., 2008) |
| 59 | W59L | (Watzka et al., 2011) |
| 59 | W59C | (Watzka et al., 2011) |
| 66 | V66M | (Oldenburg et al., 2004) |
| 66 | V66G | (Watzka et al., 2011) |
| 68 | H68Y | (Osman et al., 2006) |
| 71 | G71A | (Watzka et al., 2011) |
| 77 | N77Y | (Watzka et al., 2011) |
| 77 | N77S | (Watzka et al., 2011) |
| 123 | I123N | (Watzka et al., 2011) |
| 128 | L128R | (Oldenburg et al., 2004) |
| 139 | Y139H | (Watzka et al., 2011) |

## Additional supplementary files and source data

Supplementary Table 4. Abundance and activity data for human variants found in ClinVar, gnomAD v2 and v3, and Color Genomics dataset.

Supplementary Table 5. Reagents and resources table.

Supplementary Table 6. Names and sequences for oligos used in this paper.

Figure 1-source data 1. VKOR variant abundance and activity scores.

Figure 1-source data 2. Flow cytometry for monoclonal validation of variants.

Figure 3-source data 1. Evolutionary couplings secondary structure predictions.

Figure 3-source data 2. Evolutionary couplings 3D contact predictions.

Figure 3-source data 3. Insertion energies from Elazar et al., 2016.

Figure 5-source data 1. VKOR positional abundance and activity scores.

## METHODS

### General reagents, DNA oligonucleotides, and plasmids

Details on general reagents can be found in Supplementary Table 5. Unless otherwise noted, all chemicals were obtained from Sigma and all enzymes were obtained from New England Biolabs. *E. coli* were cultured at 37°C in Luria broth. All cell culture reagents were purchased from ThermoFisher Scientific unless otherwise noted. HEK 293T cells (ATCC CRL-3216) and derivatives thereof were cultured in Dulbecco’s modified Eagle’s medium supplemented with 10% fetal bovine serum, 100 U ml^-1^ penicillin, and 0.1 mg ml^-1^ streptomycin. Cells were induced with 2.5 ug mL^-1^ doxycycline. Cells were passaged by detachment with trypsin-EDTA 0.25%, and cells were prepared for sorting by detachment with versene. All cell lines tested negative for mycoplasma. Because our activity assay is vitamin-K dependent, all activity assays were done with the same lot of FBS to ensure similar concentrations of vitamin K in each replicate.

All synthetic oligonucleotides were obtained from IDT and can be found in Supplementary Table 6. All non-library-related plasmid modifications were performed with Gibson assembly(Gibson et al., 2009).

### Library construction

A gBLOCK with a codon-optimized sequence for human VKOR was ordered from IDT. It was then cloned into the vector pHSG298 (Clontech). Saturation mutagenesis primers were designed for each codon in VKOR from positions 2 to 163(Jain and Varadarajan, 2014) and ordered resuspended from IDT. Forward and reverse primers for each position were mixed at 2.5 mM, and used in a PCR reaction with 125 pg of pHSG298-VKOR, 5% DMSO, and 5 uL of KAPA Hifi Hotstart 2X ReadyMix. PCR products were visualized on a 0.7% agarose gel to confirm amplification of the correct product.

PCR products were then quantified using the Quant-iT PicoGreen dsDNA Assay kit (Invitrogen) using DNA control curves done in triplicate. To pool, a total amount of DNA for each reaction was calculated that maximized the volume to be drawn from the lowest concentration PCR product. Pooled PCR products were cleaned and concentrated using Zymogen Clean and Concentrate kit and then gel extracted. The pooled library was phosphorylated with T4 PNK (NEB), incubated at 37°C for 30 minutes, 65°C for 20 minutes, and then 4° indefinitely. 8.5 uL of this phosphorylated product was combined with 1 uL of 10X T4 ligase buffer (NEB) and 0.5 uL of T4 DNA ligase (NEB) to make a 10 uL overnight ligation reaction. This reaction was incubated at 16°C overnight.

The overnight ligation was then cleaned and concentrated (Zymogen) and eluted in 6 uL of ddH2O. 1 uL of this ligation was then transformed into high efficiency *E. coli* using electroporation at 2 kV. Each reaction contained 1 uL of ligation (or ligation control or pUC19 10 pg/uL) and 25 uL of E. coli. 975 uL of pre-warmed SOC media was added to each cuvette after electroporation, transferred to a culture tube, and recovered at 37°C, shaking for 1 hour. At 1 hour, 1 and 10 uL samples from all cultures were taken and plated on appropriate media (LB + kanamycin for ligation and ligation control; LB + ampicillin for pUC19), the remaining 989 uL was used to inoculate a 50 mL culture (+ kanamycin). Plates and 50 mL culture were incubated at 37°C overnight (shaking for 50 mL culture). Colonies on plates were then counted, and counts were used to calculate how many unique molecules were transformed to gauge coverage of the library. 50 mL culture was spun down and midiprepped.

To transfer the library from pKan to the recombination vector, the pKan library and recombination vector were digested with XbaI and AflII for 1 hour at 65°C. The library and cut vector were then gel extracted. The library was then ligated with the cut vector at 5:1 using T4 ligase, overnight at 16°C. The ligation was heat inactivated the next morning, clean and concentrated. Another high efficiency transformation was performed the same as described above, except this ligation was plated on LB + ampicillin (antibiotic switching strategy). Plates and 50 mL culture were incubated at 37°C overnight (shaking for 50 mL culture). Colonies on plates were then counted, and counts were used to calculate how many unique molecules had been transformed to gauge coverage of the library. A 50 mL culture was spun down and midiprepped.

To barcode individual variants, plasmid library harvested from midiprep was digested with EcoRI-HF and NdeI at 37°C for 1 hour, 65°C for 20 minutes. Barcode oligos were ordered from IDT, resuspended at 100 uM, and then annealed by combining 1 uL each of primer with 4 uL CutSmart Buffer and 34 uL ddH2O and running at 98°C for 3 minutes followed by ramping down to 25°C at −0.1°C/second. After annealing, 0.8 uL of Klenow polymerase (exonuclease negative, NEB) and 1.35 uL of 1 mM dNTPS was then combined with the 40 uL of product to fill in the barcode oligo (cycling conditions: 25°C for 15:00, 70°C for 20:00, ramp down to 37°C at −0.1°C/s). Digested vector and barcode oligo were then ligated overnight at 16°C.

The overnight ligation was then cleaned and concentrated and eluted in 6 uL of ddH2O. 1 uL of this ligation was then transformed into high efficiency *E. coli* using electroporation at 2 kV. Each reaction contained 1 uL of ligation (or ligation control or pUC19 10 pg/uL) and 25 uL of *E. coli*. 975 uL of pre-warmed SOC media was added to each cuvette after electroporation, transferred to a culture tube, and recovered at 37°C, shaking for 1 hour. At 1 hour, 1 and 10 uL samples from water and pUC19 cultures were taken and plated on LB supplemented with ampicillin. For ligation and ligation control, four flasks were prepared with 50 mLs of LB and ampicillin, and then 500 uL, 250 uL, 125 uL, 62.5 uL was sample from the 1 mL of recovery and transferred into a corresponding flask. From those flasks, 1 uL, 10 uL, and 100 uL, were sampled and plated onto LB ampicillin plates. Plates and 50 mL culture were incubated at 37°C overnight. Colonies on plates were then counted, and counts were used to calculate how many unique molecules were transformed to gauge number of barcodes. Flask with the target number of barcodes was then spun down and midiprepped.

### Cell line description

#### VAMP-seq assay cell line

HEK293T cells with a serine integrase landing pad integrated at the AAVS1 locus were used (Matreyek et al., 2017).

#### Activity assay cell line

We used a previously published reporter cell line(Haque et al., 2014) and inserted a recombinase-based landing pad at the *AAVS1* safe harbor locus using a previously published strategy (Matreyek et al., 2017). Single cell clones were transfected with TALENs for AAVS1 and the landing pad plasmid, and single cell clones were sorted. Presence of one landing pad was confirmed by 1) barcode sequencing of the landing pad and 2) co-transfection experiment with GFP and mCherry. From this, we moved forward with one clone demonstrated to have only one landing pad present (clone 45).

gRNAs to delete portions of the first exon of both *VKORC1* and *VKORC1L1* were ordered and cloned into pSpCas9(BB)-2A-GFP (PX458), which was a gift from Feng Zhang (Addgene plasmid #48138 ; http://n2t.net/addgene:48138 ; RRID:Addgene_48138). Clone 45 was then transfected with these four plasmids, and single cells were sorted based on GFP positivity. Disruption of *VKORC1* and *VKORC1L1* was confirmed by performing nested PCR, TA cloning, and then sequencing of products. We detected three alleles for both *VKORC1* and *VKORC1L1*, indicating that these loci are triploid in HEK293 cells.

A Western blot was also used to confirm absence of VKOR protein product in our activity reporter cell line. Protein lysates were harvested from ∼1 million cells using 100 uL NP40 lysis buffer with freshly prepared protease inhibitor cocktail and 1 mM PMSF. Protein lysates were Qubited for concentration, and 20 ug of each protein lysate was loaded. 4-12% BisTris NuPage gel (Thermo Fisher) was used with MES buffer + 500µl of antioxidant added to the inner chamber. The gel ran at 150V for 90 min. Gel was then transferred to Nitrocellulose using 1X transfer buffer 20% EtOH at 24V for 1 hour on ice. The blot was washed for 5 minutes with 1X TBS-T 0.1% Tween 3 times. Blot was then blocked for overnight 1X TBS-T 0.1% Tween + 5% Milk. Blot was then washed for 5 minutes with 1X TBS-T 0.1% Tween 3 times. Blot was then cut in half at the between the 25kDa and 35kDa molecular weight markers. The bottom blot was incubated with: αVKOR 1:1000 + 1X TBS-T 0.1% Tween + 5% Milk. The top blot was incubated with αbeta-actin dHRP 1:1000+ 1X TBS-T 0.1% Tween + 5% Milk. Both blots were incubated with their primary antibodies overnight at 4°C. The αVKOR blot was washed for 5 minutes with 1X TBS-T 0.1% Tween 3 times. The αVKOR blot was then incubated with 1:10,000 secondary anti-mouse-HRP (GE Healthcare NA931V) + 1X TBS-T 0.1% Tween + 5% Milk for one hour. The αbeta-actin dHRP blot remained in primary antibody during this time, as no secondary antibody is needed for a direct HRP conjugate. Both blots were then washed for 5 minutes with 1X TBS-T 0.1% Tween 3 times. Blots were then incubated with Supersignal West Dura Extended Duration Substrate (Thermo Fisher). 500µl of both substrates incubated on blot for 5 min. Blots were then dried by kimwipe and exposed using the colorimetric and chemiluminescence functions on the BioRad ChemiDoc MP (Biorad).

#### Recombination of variants in cell lines: Abundance assay

Cells were transfected in six well plates, 250,000 cells per well (12-24 wells transfected total for each experiment). Sequential transfections were performed. On day 1, 3 ug of pCAG-NLS-Bxb1 was diluted in 250 uL of OptiMEM and 6 uL of Fugene (Promega). On day 2, 3 ug of barcoded library was diluted in 250 uL of OptiMEM and 6 uL of Fugene6 and transfected. 48 hours after this second transfection, cells were induced with doxycycline at a final concentration of 2.5 ug/mL.

#### Recombination of variants in cell lines: Activity assay

Cells were transfected in six-well plates, 500,000 cells per well (18-24 wells transfected total for each experiment). 272 ng of pCAG-NLS-Bxb1 was diluted in 125 uL of OptiMEM with 2.7 ug of barcoded library. 2.25 uL of Lipofectamine 3000 (Thermo Fisher) was diluted in 125 uL of OptiMEM in a separate tube. The DNA mixture was then added to the Lipofectamine 3000 mixture and incubated at room temperature for 15 minutes. Transfection mixture was then added dropwise to one six-well plate. Cells were induced with doxycycline 48 hours after transfection, with a final concentration of 2.5 ug/mL doxycycline.

#### Enrichment sorting for recombined cells

Cells were washed once with PBS, then dissociated with versene. Media was added to dilute EDTA, and cells transferred to 15 mL conical and spun down at 300 x g for 4 minutes. Media was aspirated off, and cells were resuspended in PBS, then filtered through a 35 um nylon mesh filter. Cells were sorted on a BD Aria III FACS machine. mTagBFP2, expressed from the unrecombined landing pad, was excited with a 405 nM laser. Recombined cells either expressed mCherry (abundance) or eGFP (activity), and these were excited by a 561 nm laser and a 488 nm laser, respectively. Samples were gated for live cells using FSC-A and SSC-A, then singlets using SSC-H vs. SSC-W, FSC-H vs. FSC-W. For activity assay reporter cell line, cells were then sorted for DsRed positivity to ensure robust expression of reporter. Cells that had successfully recombined a single VKOR variant were gated on recombinant mTagBFP2 negativity and either mCherry positivity (abundance) or eGFP positivity (activity) (see Supplementary Fig. 2b for gating example). Recombined cells were sorted on “Yield” mode in the BD Diva software and grown out for 3-5 days.

#### Abundance assay quartile sorting

Recombined cells were run on a BD Aria III FACS machine. Cells were prepared for sorting as described above, and were then gated for live, recombined singlets. A ratio of eGFP/mCherry was created using the BD Diva software as a unique parameter, and the histogram of this ratio was divided into four equal bins. Each quartile was sorted into a 5 mL tube on “4-Way Purity” mode. Sorted cells were grown out for 2-4 days post sorting to ensure enough DNA for sequencing. The details of replicate sorts for activity assay are in Supplementary Table 1.

#### Antibody conjugation

Factor IX Gla domain antibody specific for carboxylation was conjugated to APC following LYNX Rapid APC Antibody Conjugation Kit instructions. Antibody was resuspended at 1 mg/mL in nuclease-free water. 1 uL of Modifier reagent was then added for every 10 uL of antibody and mixed by pipetting. That mixture was then pipetted directly onto the LYNX lyophilized mix and gently mixed by pipetting up and down twice. The conjugation mixture was then capped and incubated in the dark at room temperature overnight. After overnight incubation, 1 uL of Quencher reagent was added for every 10 uL of antibody used and left to incubate for 30 minutes. At that point, antibody was divided into 20 uL aliquots to be used for replicate experiments and stored at −20°C.

#### Activity assay antibody staining and quartile sorting

Cells were plated in six-well plates at 500,000 cells per well with D10 media with no doxycycline. All replicates were performed with 18-24 wells of cells total. After 24 hours, doxycycline was added to cells to induce expression of reporter and VKOR variant. Cells were then incubated with doxycycline for 48 hours. On day of cell sorting, each six well was washed with cold PBS, dissociated with 200 uL of versene, and then resuspended in 1 mL of phenol red-free DMEM + 1% FBS and transferred to a 5 mL FACS tube. Cells were spun at 300 x g, then washed once with 1 mL of phenol red-free DMEM + 1% FBS. Cells were spun at 300 x g, and after aspirating supernatant, cell pellet was resuspended in 100 uL of antibody diluted 1:100 in phenol red-free DMEM + 1% FBS. Cells were incubated in antibody for 20 minutes at 4°C in the dark, with vortexing at five minute intervals to ensure staining. After 20 minutes, 1 mL of staining buffer was added to each tube to dilute out antibody. Cells were spun at 300 x g, washed twice more similarly with staining buffer, then resuspended in 200 uL. At this point, all tubes were pooled and filtered to remove clumps. Cells were then sorted using a FACSAria III (BD Biosciences) into bins based on their APC intensity. First, live, single, recombinant cells were selected as described above. A histogram of APC was created and gates dividing the library into four equally populated bins based on APC fluorescence intensity were drawn. The details of replicate sorts for activity assay are in Supplementary Table 2.

### gDNA prep, barcode amplification, and sequencing

Cells were then collected, pelleted by centrifugation and stored at −20 °C. Genomic DNA was prepared using a DNEasy kit, according to the manufacturer’s instructions (Qiagen), with the addition of a 30 min incubation at 37 °C with RNAse in the re-suspension step. Eight 50 μl first-round PCR reactions were each prepared with a final concentration of ∼50 ng μl−1 input genomic DNA, 1 × Q5 High-Fidelity Master Mix and 0.25 μM of the KAM499/VKORampR 1.1 primers. The reaction conditions were 98 °C for 30 s, 98°C for 10 s, 65°C for 20 s, 72°C for 60 s, repeat 5 times, 72°C for 2 min, 4°C hold. Eight 50 μl reactions were combined, bound to AMPure XP (Beckman Coulter), cleaned and eluted with 21 μl water. Forty percent of the eluted volume was mixed with Q5 High-Fidelity Master Mix; VKOR_indexF_1.1 and one of the indexed reverse primers, PTEN_seq_R1a through PTEN_seq_R2a, were added at 0.25 μM each. These reactions were run with Sybr Green I on a BioRad MiniOpticon; reactions were denatured for 3 minutes at 95°C and cycled 20 times at 95°C for 15s, 60°C for 15s, 72°C for 15s with a final 3 min extension at 72°C. The indexed amplicons were mixed based in relative fluorescence units and run on a 1% agarose gel with Sybr Safe and gel extracted using a freeze and squeeze column (Bio-Rad). The product was quantified using Kapa Illumina Quant kit.

### Subassembly

Barcoded VKOR library was subassembled using a MiSeq 600 kit (Illumina). Two amplicons were generated, one forward, one reverse. PCR reactions were each prepared with ∼500 ng input plasmid DNA, 1 × KAPA High-Fidelity Master Mix and 0.25 μM of the VKOR_SA_amp_F/VKOR_SA_amp_R or VKOR_SA_for_amp_R2.0/VKOR_SA_rev_amp_F2.0 primers. PCR reactions were run at 95°C for five minutes, then cycled 15 times at 98°C for 0:20, 60°C for 0:15, 72°C for 0:30, with a final extension at 72°C for 2:00. Amplicons (741 bp) were gel extracted on a 1.0% gel run at 130V for 35 mins. The product was quantified using Qubit and Kapa Illumina quant kit. Read lengths were as follows: 289 bp forward read, 18 bp index1, 18 bp index 2 (index = barcode forward and reverse). All reads sharing a common barcode sequence were collapsed to form the consensus variant sequence, resulting in 175,052 barcodes after filtering.

### Barcode counting and variant calling

Enrich2 was used to quantify barcodes from bin sequencing, using a minimum quality filter of 20 (Rubin et al., 2017). FASTQ files containing barcodes and the barcode map for VKOR were used as input for Enrich2. Enrich2 configuration files for each experiment are available on the GitHub repository. Barcodes assigned to variants containing insertion, deletions, or multiple amino-acid alterations were removed from the analysis.

### Calculating scores and classifications

Scores and classifications were assigned using previously published analysis pipeline (Matreyek et al., 2018). Briefly, for each protein variant, frequencies in each bin were calculated by dividing counts by total counts. From there, we filtered variants based on the number of experiments in which it was observed (F_expt_ = 2) and their frequency (F_freq_ = 10^-4^), after noticing that low frequency variants introduced noise to the analysis. These frequencies were then each weighted by multiplying by 0.25, 0.5, 0.75, and 1 in a bin-wise fashion. We generated a replicate score for each variant by using min-max normalization: normalizing to the median weighted average of the nonsense distribution set at 0 and the median weighted average of the synonymous distribution set at 1. We then averaged those scores for a final, experiment-wide variant score. Standard deviation and standard error were also calculated for each variant, and 95% confidence intervals were estimated using standard error, assuming a normal distribution. Abundance and activity classifications were assigned by assessing variant score and confidence intervals in relation to synonymous variant distribution. To do this, we established a cut-off that separated the 5% of synonymous variants with the lowest abundance (or activity) scores from the 95% of synonymous variants with higher abundance (or activity) scores. Variants with both scores and upper confidence intervals below this threshold were classified as “low,” while those with scores below but upper confidence above were classified as “possibly low.” Variants with scores and lower confidence intervals above the threshold were classified as “WT-like”, while those with scores above lower confidence intervals below the threshold were classified as “possibly WT-like.” Finally, another threshold was set that separated the 5% of synonymous variants with the highest scores from the rest of the synonymous distribution. Variants that had scores above this threshold, with lower confidence intervals above the lower threshold were classified as “high.”

### Windowed abundance and activity analysis

Windowed averages of abundance and activity scores were calculated using a window length of 10 positions with center alignment. Scores were calculated for both charged amino acids (R, K, H, D, E,) and aliphatic amino acids (G, A, V, I, L).

### Evolutionary couplings analysis

EVcouplings extracts the constraints between pairs of residues, as evidenced in alignments of homologous sequences: first homologous sequences must be collected and aligned, and then a model of statistical energy costs and benefits between residues is fit to explain the sequence variation in the alignment. We collected an alignment of 2770 sequences using jackhammer (http://hmmer.org/) to query the human VKOR sequence against UniRef100 (https://www.uniprot.org/uniref/), with a bitscore per residue cutoffs of 0.4 and 7 search iterations. We predicted secondary structures where the summed strength of couplings at would-be alpha helix and beta strand contacts scored above 1.5 for alpha helices and 0.75 respectively, for two or more consecutive residues. We extended the called helices and strands by one residue on each side for a minimum structure size of four residues. All methods used for building alignments, training the model, folding, and predicting secondary structure are part of the EVcouplings software (https://evcouplings.org/) (Hopf et al., 2019).

### Homology modeling

A homology model of human VKOR was made by accessing I-TASSER (Yang et al., 2015) and using PDB structure 4NV5 as a template for threading. Model1 from results was used for all figures in this paper.

### Hierarchical clustering

Hierarchical clustering was performed on abundance score vectors for each position using the hclust function in R. Dendrogram for hierarchically clustered heatmap was drawn using dendextend package (version 1.12.0).

### Active site residue analysis

Activity and abundance scores were rescaled so that the lowest score present in the dataset was set at 0, and the highest score at 1. A ratio of rescaled activity to rescaled abundance (specific activity) was then calculated for every variant. Using variant specific activity scores, median specific activity was calculated for each position. Threshold for classification as an active site position was drawn based on scores of known redox cysteines at positions 132 and 135, resulting in lowest 12.5% of median specific activity scores being classified as active site residues. We additionally required that any position within this group had been scored for at least four variants to eliminate noise from poor sampling.

### Data availability

All raw sequence data and function scores are freely available for all academic users by non-exclusive license under reasonable terms to commercial entities that have committed to open sharing of VKOR sequence variants and under a free non-exclusive license to non-profit entities. The Illumina raw sequencing files and barcode–variant maps can be accessed at the NCBI Gene Expression Omnibus (GEO) repository under accession number GSE149922. The data presented in the manuscript are available as Supplementary Data files.

### Code availability

Code for analysis is available at http://github.com/FowlerLab/VKOR. The code used to train the evolutionary couplings model is available at the EVcouplings GitHub repository (https://github.com/debbiemarkslab/EVcouplings). The data used to train the model is publically available at uniprot (https://uniprot.org).

## ACKNOWLEDGEMENTS

We thank D. Nickerson and M. Dunham for helpful conversations in analyzing the data and writing the manuscript, A. Leith of the UW Foege Flow Lab and D. Prunkard of the UW Pathology Flow Cytometry Core Facility for assistance with cell sorting, and all members of the Fowler lab for helpful feedback on figures. This work was supported by the NIH (R24GM115277 P01GM116691), by the Canada Excellence Research Chairs Program, and by a Canadian Institutes of Health Research Foundation Award. M.A.C. was supported by a National Science Foundation Graduate Research Fellowship. D.M. and N.J.R. thank the Chan Zuckerberg Initiative DAF (grant number 2018-191853) for financial support.

## AUTHOR CONTRIBUTIONS

M.A.C. carried out abundance and activity experiments and analyzed the data. N.R. and D.M. performed evolutionary couplings analysis. J.S. prepared samples for next-generation sequencing. K.S. cloned variants for abundance assay validation. K.A.M. provided abundance analysis framework. D.D. provided human *VKORC1* variants from Color Genomics. A.R. provided the parental activity reporter cell line. M.V., S.S. and F.P.R. provided a mutagenized VKOR library. M.A.C. and D.M.F. wrote the manuscript with input from co-authors.

## REFERENCES

Bodin L, Perdu J, Diry M, Horellou M-H, Loriot M-A. 2008. Multiple genetic alterations in vitamin K epoxide reductase complex subunit 1 gene (VKORC1) can explain the high dose requirement during oral anticoagulation in humans. J Thromb Haemost 6:1436–1439. doi:10.1111/j.1538-7836.2008.03049.x

Czogalla KJ, Biswas A, Höning K, Hornung V, Liphardt K, Watzka M, Oldenburg J. 2016. Warfarin and vitamin K compete for binding to Phe55 in human VKOR. Nat Struct Mol Biol. doi:10.1038/nsmb.3338

Czogalla KJ, Biswas A, Rost S, Watzka M, Oldenburg J. 2014. The Arg98Trp mutation in human VKORC1 causing VKCFD2 disrupts a di-Arginine-based ER retention motif. Blood. doi:10.1182/blood-2013-12-545988

Elazar AA, Weinstein J, Biran I, Fridman Y, Bibi E, Fleishman SJ. 2016. Mutational scanning reveals the determinants of protein insertion and association energetics in the plasma membrane. eLife Sciences e12125. doi:10.7554/eLife.12125

Elazar A, Weinstein JJ, Prilusky J, Fleishman SJ. 2016. Interplay between hydrophobicity and the positive-inside rule in determining membrane-protein topology. Proc Natl Acad Sci U S A 113:10340–10345. doi:10.1073/pnas.1605888113

Fleming KG, Engelman DM. 2001. Specificity in transmembrane helix-helix interactions can define a hierarchy of stability for sequence variants. Proc Natl Acad Sci U S A 98:14340– 14344. doi:10.1073/pnas.251367498

Gasperini M, Starita L, Shendure J. 2016. The power of multiplexed functional analysis of genetic variants. Nat Protoc 11:1782–1787. doi:10.1038/nprot.2016.135

Gibson DG, Young L, Chuang R-Y, Venter JC, Hutchison CA 3rd, Smith HO. 2009. Enzymatic assembly of DNA molecules up to several hundred kilobases. Nat Methods 6:343–345. doi:10.1038/nmeth.1318

Gong IY, Schwarz UI, Crown N, Dresser GK, Lazo-Langner A, Zou G, Roden DM, Stein CM, Rodger M, Wells PS, Kim RB, Tirona RG. 2011. Clinical and genetic determinants of warfarin pharmacokinetics and pharmacodynamics during treatment initiation. PLoS One 6:e27808. doi:10.1371/journal.pone.0027808

Gray VE, Hause RJ, Fowler DM. 2017. Analysis of Large-Scale Mutagenesis Data To Assess the Impact of Single Amino Acid Substitutions. Genetics 207:53–61. doi:10.1534/genetics.117.300064

Guerriero CJ, Reutter K-R, Augustine AA, Preston GM, Weiberth KF, Mackie TD, Cleveland-Rubeor HC, Bethel NP, Callenberg KM, Nakatsukasa K, Grabe M, Brodsky JL. 2017. Transmembrane helix hydrophobicity is an energetic barrier during the retrotranslocation of integral membrane ERAD substrates. Mol Biol Cell 28:2076–2090. doi:10.1091/mbc.E17-03-0184

Hallgren KW, Qian W, Yakubenko AV, Runge KW, Berkner KL. 2006. r-VKORC1 Expression in Factor IX BHK Cells Increases the Extent of Factor IX Carboxylation but Is Limited by Saturation of Another Carboxylation Component or by a Shift in the Rate-Limiting Step†. Biochemistry 45:5587–5598. doi:10.1021/bi051986y

Haque JA, McDonald MG, Kulman JD, Rettie AE. 2014. A cellular system for quantitation of vitamin K cycle activity: structure-activity effects on vitamin K antagonism by warfarin metabolites. Blood 123:582–589. doi:10.1182/blood-2013-05-505123

Harrington DJ, Gorska R, Wheeler R, Davidson S, Murden S, Morse C, Shearer MJ, Mumford AD. 2008. Pharmacodynamic resistance to warfarin is associated with nucleotide substitutions in VKORC1. J Thromb Haemost 6:1663–1670. doi:10.1111/j.1538-7836.2008.03116.x

Harrington DJ, Siddiq S, Allford SL, Shearer MJ, Mumford AD. 2011. More on: endoplasmic reticulum loop VKORC1 substitutions cause warfarin resistance but do not diminish gamma-carboxylation of the vitamin K-dependent coagulation factors. J Thromb Haemost 9:1093–1095. doi:10.1111/j.1538-7836.2011.04249.x

Hatahet F, Blazyk JL, Martineau E, Mandela E, Zhao Y, Campbell RE, Beckwith J, Boyd D. 2015. Altered Escherichia coli membrane protein assembly machinery allows proper membrane assembly of eukaryotic protein vitamin K epoxide reductase. Proc Natl Acad Sci U S A 112:15184–15189. doi:10.1073/pnas.1521260112

Hessa T, Meindl-Beinker NM, Bernsel A, Kim H, Sato Y, Lerch-Bader M, Nilsson I, White SH, von Heijne G. 2007. Molecular code for transmembrane-helix recognition by the Sec61 translocon. Nature 450:1026–1030. doi:10.1038/nature06387

Hessa T, Sharma A, Mariappan M, Eshleman HD, Gutierrez E, Hegde RS. 2011. Protein targeting and degradation are coupled for elimination of mislocalized proteins. Nature 475:394–397. doi:10.1038/nature10181

Hopf TA, Colwell LJ, Sheridan R, Rost B, Sander C, Marks DS. 2012. Three-Dimensional Structures of Membrane Proteins from Genomic Sequencing. Cell 149:1607–1621. doi:10.1016/j.cell.2012.04.012

Hopf TA, Green AG, Schubert B, Mersmann S, Schärfe CPI, Ingraham JB, Toth-Petroczy A, Brock K, Riesselman AJ, Palmedo P, Kang C, Sheridan R, Draizen EJ, Dallago C, Sander C, Marks DS. 2019. The EVcouplings Python framework for coevolutionary sequence analysis. Bioinformatics 35:1582–1584. doi:10.1093/bioinformatics/bty862

Jain PC, Varadarajan R. 2014. A rapid, efficient, and economical inverse polymerase chain reaction-based method for generating a site saturation mutant library. Anal Biochem 449:90–98. doi:10.1016/j.ab.2013.12.002

Kabsch W, Sander C. 1983. Dictionary of protein secondary structure: pattern recognition of hydrogen-bonded and geometrical features. Biopolymers 22:2577–2637. doi:10.1002/bip.360221211

Karczewski KJ, Francioli LC, Tiao G, Cummings BB, Alföldi J, Wang Q, Collins RL, Laricchia KM, Ganna A, Birnbaum DP, Gauthier LD, Brand H, Solomonson M, Watts NA, Rhodes D, Singer-Berk M, England EM, Seaby EG, Kosmicki JA, Walters RK, Tashman K, Farjoun Y, Banks E, Poterba T, Wang A, Seed C, Whiffin N, Chong JX, Samocha KE, Pierce-Hoffman E, Zappala Z, O’Donnell-Luria AH, Minikel EV, Weisburd B, Lek M, Ware JS, Vittal C, Armean IM, Bergelson L, Cibulskis K, Connolly KM, Covarrubias M, Donnelly S, Ferriera S, Gabriel S, Gentry J, Gupta N, Jeandet T, Kaplan D, Llanwarne C, Munshi R, Novod S, Petrillo N, Roazen D, Ruano-Rubio V, Saltzman A, Schleicher M, Soto J, Tibbetts K, Tolonen C, Wade G, Talkowski ME, The Genome Aggregation Database Consortium, Neale BM, Daly MJ, MacArthur DG. 2019. Variation across 141,456 human exomes and genomes reveals the spectrum of loss-of-function intolerance across human protein-coding genes. bioRxiv. doi:10.1101/531210

Kurnik D, Qasim H, Sominsky S, Lubetsky A, Markovits N, Li C, Stein CM, Halkin H, Gak E, Loebstein R. 2012. Effect of the VKORC1 D36Y variant on warfarin dose requirement and pharmacogenetic dose prediction. Thromb Haemost 108:781–788. doi:10.1160/TH12-03-0151

Landrum MJ, Lee JM, Riley GR, Jang W, Rubinstein WS, Church DM, Maglott DR. 2014. ClinVar: public archive of relationships among sequence variation and human phenotype. Nucleic Acids Res 42:D980–5. doi:10.1093/nar/gkt1113

Li T, Chang C-Y, Jin D-Y, Lin P-J, Khvorova A, Stafford DW. 2004. Identification of the gene for vitamin K epoxide reductase. Nature 427:541–544. doi:10.1038/nature02254

Liu S, Cheng W, Fowle Grider R, Shen G, Li W. 2014. Structures of an intramembrane vitamin K epoxide reductase homolog reveal control mechanisms for electron transfer. Nat Commun 5. doi:10.1038/ncomms4110

Li W, Schulman S, Dutton RJ, Boyd D, Beckwith J, Rapoport TA. 2010. Structure of a bacterial homologue of vitamin K epoxide reductase. Nature 463:507–512. doi:10.1038/nature08720

Loebstein R, Dvoskin I, Halkin H, Vecsler M, Lubetsky A, Rechavi G, Amariglio N, Cohen Y, Ken-Dror G, Almog S, Gak E. 2007. A coding VKORC1 Asp36Tyr polymorphism predisposes to warfarin resistance. Blood 109:2477–2480. doi:10.1182/blood-2006-08-038984

Marks DS, Colwell LJ, Sheridan R, Hopf TA, Pagnani A, Zecchina R, Sander C. 2011. Protein 3D structure computed from evolutionary sequence variation. PLoS One 6:e28766. doi:10.1371/journal.pone.0028766

Matreyek KA, Starita LM, Stephany JJ, Martin B, Chiasson MA, Gray VE, Kircher M, Khechaduri A, Dines JN, Hause RJ, Bhatia S, Evans WE, Relling MV, Yang W, Shendure J, Fowler DM. 2018. Multiplex assessment of protein variant abundance by massively parallel sequencing. Nat Genet 50:874–882. doi:10.1038/s41588-018-0122-z

Matreyek KA, Stephany JJ, Fowler DM. 2017. A platform for functional assessment of large variant libraries in mammalian cells. Nucleic Acids Res. doi:10.1093/nar/gkx183

Ma W, Goldberg J. 2013. Rules for the recognition of dilysine retrieval motifs by coatomer. EMBO J 32:926–937. doi:10.1038/emboj.2013.41

Merkle PS, Gotfryd K, Cuendet MA, Leth-Espensen KZ, Gether U, Loland CJ, Rand KD. 2018. Substrate-modulated unwinding of transmembrane helices in the NSS transporter LeuT. Sci Adv 4:eaar6179. doi:10.1126/sciadv.aar6179

Mravic M, Thomaston JL, Tucker M, Solomon PE, Liu L, DeGrado WF. 2019. Packing of apolar side chains enables accurate design of highly stable membrane proteins. Science 363:1418–1423. doi:10.1126/science.aav7541

Nilsson I, von Heijne G. 1990. Fine-tuning the topology of a polytopic membrane protein: role of positively and negatively charged amino acids. Cell 62:1135–1141. doi:10.1016/0092-8674(90)90390-z

Oldenburg J, Rost S, Fregin A, Geisen C, Ivaskevicius V, Seifried E, Scharrer I, Heistinger M, Tuddenham E, Muller-Reible C, Zieger B. 2004. Mutations in the VKORC1 Gene Cause Warfarin Resistance, Warfarin Sensitivity and Combined Deficiency of Vitamin K Dependent Coagulation Factors. Blood 104:277–277.

Osinbowale O, Al Malki M, Schade A, Bartholomew JR. 2009. An algorithm for managing warfarin resistance. Cleve Clin J Med 76:724–730. doi:10.3949/ccjm.76a.09062

Osman A, Enström C, Arbring K, Söderkvist P, Lindahl TL. 2006. Main haplotypes and mutational analysis of vitamin K epoxide reductase (VKORC1) in a Swedish population: a retrospective analysis of case records. J Thromb Haemost 4:1723–1729. doi:10.1111/j.1538-7836.2006.02039.x

Owen RP, Gong L, Sagreiya H, Klein TE, Altman RB. 2010. VKORC1 Pharmacogenomics Summary. Pharmacogenet Genomics 20:642–644. doi:10.1097/FPC.0b013e32833433b6

Peoc’h K, Pruvot S, Gourmel C, dit Sollier CB, Drouet L. 2009. A new VKORC1 mutation leading to an isolated resistance to fluindione. Br J Haematol 145:841–843. doi:10.1111/j.1365-2141.2009.07687.x

Rieder MJ, Reiner AP, Gage BF, Nickerson DA, Eby CS, McLeod HL, Blough DK, Thummel KE, Veenstra DL, Rettie AE. 2005. Effect of VKORC1 Haplotypes on Transcriptional Regulation and Warfarin Dose. N Engl J Med 352:2285–2293. doi:10.1056/NEJMoa044503

Rishavy MA, Usubalieva A, Hallgren KW, Berkner KL. 2011. Novel Insight into the Mechanism of the Vitamin K Oxidoreductase (VKOR) ELECTRON RELAY THROUGH Cys43 AND Cys51 REDUCES VKOR TO ALLOW VITAMIN K REDUCTION AND FACILITATION OF VITAMIN K-DEPENDENT PROTEIN CARBOXYLATION. J Biol Chem 286:7267–7278. doi:10.1074/jbc.M110.172213

Rost S, Fregin A, Ivaskevicius V, Conzelmann E, Hörtnagel K, Pelz H-J, Lappegard K, Seifried E, Scharrer I, Tuddenham EGD, Müller CR, Strom TM, Oldenburg J. 2004. Mutations in VKORC1 cause warfarin resistance and multiple coagulation factor deficiency type 2. Nature 427:537–541. doi:10.1038/nature02214

Rubin AF, Gelman H, Lucas N, Bajjalieh SM, Papenfuss AT, Speed TP, Fowler DM. 2017. A statistical framework for analyzing deep mutational scanning data. Genome Biol 18:150. doi:10.1186/s13059-017-1272-5

Schmeits PCJ, Hermans MHA, van Geest-Daalderop JHH, Poodt J, de Sauvage Nolting PRW, Conemans JMH. 2010. VKORC1 mutations in patients with partial resistance to phenprocoumon. Br J Haematol 148:955–957. doi:10.1111/j.1365-2141.2009.08017.x

Schulman S, Wang B, Li W, Rapoport TA. 2010. Vitamin K epoxide reductase prefers ER membrane-anchored thioredoxin-like redox partners. Proc Natl Acad Sci U S A 107:15027– 15032. doi:10.1073/pnas.1009972107

Sharpe HJ, Stevens TJ, Munro S. 2010. A comprehensive comparison of transmembrane domains reveals organelle-specific properties. Cell 142:158–169. doi:10.1016/j.cell.2010.05.037

Shen G, Cui W, Zhang H, Zhou F, Huang W, Liu Q, Yang Y, Li S, Bowman GR, Sadler JE, Gross ML, Li W. 2017. Warfarin traps human vitamin K epoxide reductase in an intermediate state during electron transfer. Nat Struct Mol Biol 24:69. doi:10.1038/nsmb.3333

Tie J-K, Jin D-Y, Stafford DW. 2012. Human Vitamin K Epoxide Reductase and Its Bacterial Homologue Have Different Membrane Topologies and Reaction Mechanisms. J Biol Chem 287:33945–33955. doi:10.1074/jbc.M112.402941

Tie J-K, Jin D-Y, Tie K, Stafford DW. 2013. Evaluation of warfarin resistance using transcription activator-like effector nucleases-mediated vitamin K epoxide reductase knockout HEK293 cells. J Thromb Haemost 11:1556–1564. doi:10.1111/jth.12306

Toth-Petroczy A, Palmedo P, Ingraham J, Hopf TA, Berger B, Sander C, Marks DS. 2016. Structured States of Disordered Proteins from Genomic Sequences. Cell 167:158–170.e12. doi:10.1016/j.cell.2016.09.010

Touw WG, Baakman C, Black J, te Beek TAH, Krieger E, Joosten RP, Vriend G. 2015. A series of PDB-related databanks for everyday needs. Nucleic Acids Res 43:D364–8. doi:10.1093/nar/gku1028

Ulmschneider MB, Sansom MS. 2001. Amino acid distributions in integral membrane protein structures. Biochim Biophys Acta 1512:1–14. doi:10.1016/s0005-2736(01)00299-1

von Heijne G. 1989. Control of topology and mode of assembly of a polytopic membrane protein by positively charged residues. Nature 341:456–458. doi:10.1038/341456a0

Watzka M, Geisen C, Bevans CG, Sittinger K, Spohn G, Rost S, Seifried E, Müller CR, Oldenburg J. 2011. Thirteen novel VKORC1 mutations associated with oral anticoagulant resistance: insights into improved patient diagnosis and treatment. J Thromb Haemost 9:109–118. doi:10.1111/j.1538-7836.2010.04095.x

Wilms EB, Touw DJ, Conemans JMH, Veldkamp R, Hermans M. 2008. A new VKORC1 allelic variant (p.Trp59Arg) in a patient with partial resistance to acenocoumarol and phenprocoumon. J Thromb Haemost 6:1224–1226. doi:10.1111/j.1538-7836.2008.02975.x

Wu S, Chen X, Jin D-Y, Stafford DW, Pedersen LG, Tie J-K. 2018. Warfarin and vitamin K epoxide reductase: a molecular accounting for observed inhibition. Blood. doi:10.1182/blood-2018-01-830901

Yang J, Yan R, Roy A, Xu D, Poisson J, Zhang Y. 2015. The I-TASSER Suite: protein structure and function prediction. Nat Methods 12:7–8. doi:10.1038/nmeth.3213

Yuan H-Y, Chen J-J, Lee MTM, Wung J-C, Chen Y-F, Charng M-J, Lu M-J, Hung C-R, Wei C-Y, Chen C-H, Wu J-Y, Chen Y-T. 2005. A novel functional VKORC1 promoter polymorphism is associated with inter-individual and inter-ethnic differences in warfarin sensitivity. Hum Mol Genet 14:1745–1751. doi:10.1093/hmg/ddi180

Zimmermann A, Matschiner JT. 1974. Biochemical basis of hereditary resistance to warfarin in the rat. Biochem Pharmacol 23:1033–1040. doi:10.1016/0006-2952(74)90002-1

